# Mammalian Cell Proliferation Requires Noncatalytic Functions of O-GlcNAc Transferase

**DOI:** 10.1101/2020.10.22.351288

**Authors:** Zebulon G. Levine, Sarah C. Potter, Cassandra M. Joiner, George Q. Fei, Behnam Nabet, Matthew Sonnett, Natasha E. Zachara, Nathanael S. Gray, Joao A. Paulo, Suzanne Walker

## Abstract

O-GlcNAc transferase (OGT), found in the nucleus and cytoplasm of all mammalian cell types, is essential for cell proliferation. Why OGT is required for cell growth is not known. OGT performs two enzymatic reactions in the same active site. In one, it glycosylates thousands of different proteins, and in the other, it proteolytically cleaves another essential protein involved in gene expression. Deconvoluting OGT’s myriad cellular roles has been challenging because genetic deletion is lethal; complementation methods have not been established. Here, we developed approaches to replace endogenous OGT with separation-of-function variants to investigate the importance of OGT’s enzymatic activities for cell viability. Using genetic complementation, we found that OGT’s glycosyltransferase function is required for cell growth but its protease function is dispensable. We next used complementation to construct a cell line with degron-tagged wild-type OGT. When OGT was degraded to very low levels, cells stopped proliferating but remained viable. Adding back catalytically-inactive OGT rescued growth. Therefore, OGT has an essential noncatalytic role that is necessary for cell proliferation. By developing a method to quantify how OGT’s catalytic and noncatalytic activities affect protein abundance, we found that OGT’s noncatalytic functions often affect different proteins from its catalytic functions. Proteins involved in oxidative phosphorylation and the actin cytoskeleton were especially impacted by the noncatalytic functions. We conclude that OGT integrates both catalytic and noncatalytic functions to control cell physiology.

**Significance:** Mammalian cells contain only one glycosyltransferase, OGT, that operates in the nucleus and cytoplasm rather than the secretory pathway. OGT is required for cell proliferation, but a basic unanswered question is what OGT functions are essential. This question is challenging to address because OGT has thousands of glycosylation substrates, two different enzymatic activities, and a large number of binding partners. Here, by establishing genetic tools to replace endogenous OGT with variants that preserve only a subset of its activities, we show that only a low level of glycosylation activity is required to maintain cell viability; however, cell proliferation requires noncatalytic OGT function(s). The ability to replace OGT with variants provides a path to identifying its essential substrates and binding partners.

## Introduction

O-linked *N*-acetylglucosamine transferase (OGT) is the most conserved glycosyltransferase encoded in the human genome (Figure 1D, S1A) (1, 2) and is essential for mammalian cell survival (3–5). Unlike other glycosyltransferases, which act in the endoplasmic reticulum and Golgi apparatus, OGT is found in the nucleus, cytoplasm, and mitochondria (6, 7). OGT attaches the monosaccharide O-linked N-acetylglucosamine (O-GlcNAc) to serine and threonine side chains of thousands of proteins (Figure 1A) (6, 8). These O-GlcNAc modifications are often dynamic and can be removed by the glycosidase O-GlcNAcase (OGA) (6, 9, 10). Because protein O-GlcNAc levels are sensitive to nutrient conditions and are cytoprotective against multiple forms of cellular stress, the modification is thought to be important in maintaining cellular homeostasis (6, 11–13). High O-GlcNAc levels correlate with aggressiveness for multiple forms of cancer (14–17) and have also been implicated in the pathogenesis of metabolic syndrome. It is therefore speculated that inhibiting OGT’s catalytic activity will lead to therapeutic benefit for treating cancer or cardiometabolic disease (8, 18–21).

**Figure 1:**
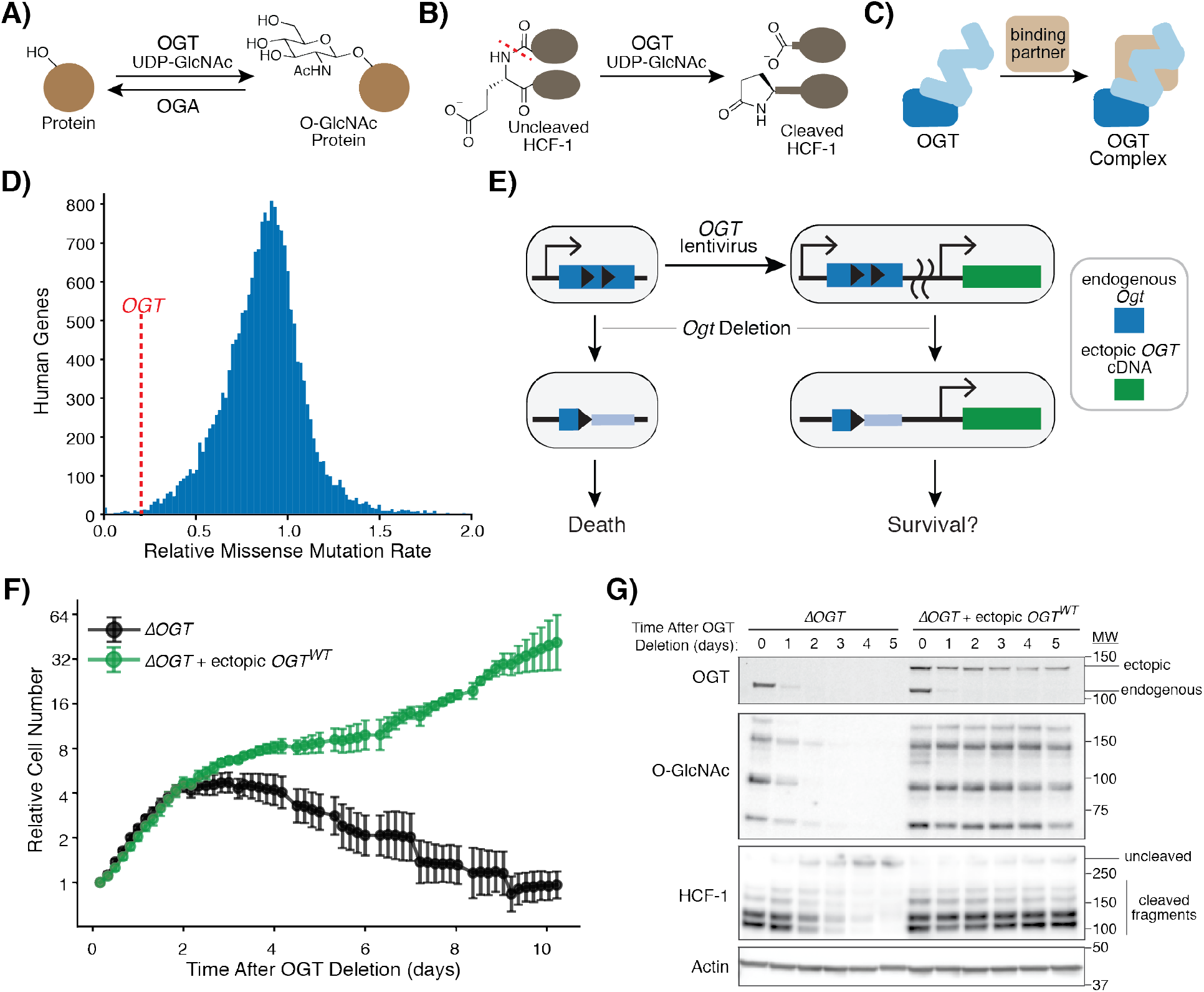
Ectopic Expression of OGT Complements Loss of Endogenous Gene. **A)** OGT transfers GlcNAc from UDP-GlcNAc to Ser or Thr residues of many nuclear and cytoplasmic proteins. OGA removes O-GlcNAc. **B)** OGT cleaves essential transcriptional regulator HCF-1. Glutamate glycosylation induces peptide backbone rearrangement and cleavage, which leaves a pyroglutamate on the C-terminal fragment. **C)** OGT forms multiple protein complexes. **D)** *OGT* is one of the most conserved human genes. Histogram shows genome-wide amino acid conservation based on a human exome database (2). *OGT* is marked by the dashed red line at 0.20. X-axis is truncated at 2. **E)** *OGT* replacement at an ectopic locus can test which OGT features are required for cell survival. Engineered MEFs (40) allow deletion of endogenous *Ogt*. We tested if lentiviral introduction of a single isoform of *OGT* could complement defects in cell survival and OGT activity. **F)** Ectopic *OGT* expression rescues cell growth. Cells with or without ectopic *OGT* were imaged every ~4 hours after inducing *Ogt* knockout at t=0. Relative cell number is based on normalized measurements of confluence from live cell imaging. Error bars are SD (n = 6 per condition). **G)** Ectopic *OGT* expression rescues biochemical activities lost upon *Ogt* knockout. Immunoblot analysis shows endogenous and ectopic OGT, protein O-GlcNAc bands (measured by pan-O-GlcNAc monoclonal antibody RL2), and uncleaved HCF-1 or cleaved fragments. See also Figure S1A–F.

OGT is essential for cell viability and impacts numerous aspects of cell physiology including metabolism, gene expression, and cell signaling. Its broad effects have been attributed to its widespread Ser/Thr glycosylation activity (8, 22, 23), yet OGT has other biochemical functions that may be important for viability. In an unusual example of a physiologically-relevant second enzymatic activity that occurs in the same active site, OGT proteolytically cleaves the essential transcriptional co-regulator HCF-1 (Figure 1B, S1B) (24–26); this proteolytic maturation process is proposed be critical for cytokinesis (26–29). In addition to catalyzing both glycosylation and cleavage reactions, OGT interacts with binding partners (Figure 1C) that recruit OGT to specific substrates (3, 30–34). Recent evidence suggests OGT may regulate *C. elegans* physiology independent of its catalytic function (36, 37); however, given that OGT is not essential in *C. elegans* (38), it remains unclear whether OGT’s binding interactions play major roles in regulating mammalian physiology independent of OGT’s enzymatic activity (35, 39).

Here we developed a complementation strategy to evaluate whether OGT’s catalytic functions are essential for mammalian cell growth. We then modified the strategy to evaluate contributions of OGT’s noncatalytic functions to cell proliferation and to link OGT’s biochemical activities to pathways they control. Our study answers the fundamental question of which of OGT’s biochemical activities are needed for cell growth, has implications for how to target OGT in cancer and metabolic disease, and provides tools and approaches that will lead to a better understanding of this unusual protein.

## Results

### Inducible *OGT* complementation rescues cell growth

To test which OGT functions are required for cell survival, we developed a complementation system starting from a mouse embryonic fibroblast (MEF) cell line in which Cre-recombinase-mediated *Ogt* knockout is induced by drug treatment (4, 5, 40). Because OGT is normally produced as multiple isoforms (8), we first tested whether a single isoform could rescue knockout of endogenous *Ogt* (Figure 1E). We infected MEFs with a lentiviral vector (41) encoding human OGT, which has 99.8% amino acid identity with that of mouse OGT. Human OGT was produced as a fusion to the red fluorescent protein mKate2; FACS-based selection of cells with sufficient ectopic OGT levels corrected for variable transgene expression due to stochastic lentiviral integration into the cell genome (Figure S1C–D) (42, 43). To test for complementation, we tracked cell abundance by live cell imaging after generating an endogenous *Ogt* knockout. As expected (40), cells without *Ogt* underwent growth arrest at two days and died over the subsequent week (Figures 1F, S1E). Cells lacking endogenous *Ogt* but harboring an ectopic copy of human *OGT* continued to grow, showing that a single OGT isoform can rescue growth (Figure 1F).

Immunoblot analysis of relevant biomarkers confirmed that genetic complementation maintained OGT’s catalytic activities (Figure 1G). OGT produced from the endogenous gene (Figure 1G, OGT, lower band) became undetectable two days after knockout, and in the absence of ectopic *OGT*, we observed loss of protein O-GlcNAc bands and accumulation of uncleaved HCF-1. OGA synthesis, which is downregulated by low protein O-GlcNAc levels (40, 44),greatly decreased without OGT (Figure S1F). By day 3, we observed a 42 kDa PARP-1 fragment associated with necrosis (Figure S1F) (45, 46). However, in cells producing mKate2-OGT, protein O-GlcNAc levels and HCF-1 cleavage remained high after *Ogt* knockout, and accumulation of the necrosis-associated PARP-1 fragment was minimal. Cells expressing untagged ectopic *OGT,* which could not be selected for sufficient protein expression, had high amounts of the necrosis-associated PARP-1 fragment by day 3 (Figure S1F), consistent with OGT levels being insufficient to rescue cell growth (Figures S1E–F). Taken together, our results show that OGT’s canonical 1036 amino acid isoform, when produced at sufficient levels, complements both cell growth and OGT’s catalytic activities. Therefore, this single isoform encodes all of OGT’s essential functions. A key advantage to ectopic expression of an *OGT* cDNA from a constitutive promoter is that O-GlcNAc-responsive mRNA regulation is bypassed, facilitating comparison of different OGT variants to identify those that support survival (44, 47–49).

### Ser/ Thr Glycosylation is OGT’ s Only Essential Catalytic Activity

We tested a set of previously characterized OGT variants in our complementation system to determine which catalytic activities were necessary for cell survival (Figure 2) (24, 25, 50–54). Ser/Thr glycosyation and HCF-1 cleavage occur in the same active site, and both reactions use UDP-GlcNAc as a substrate (24, 51, 52). Active site residue K842 enables GlcNAc transfer for both reactions, so mutating this residue renders OGT catalytically inactive (24, 52, 54). An OGT^K842M^ variant was used to assess whether OGT has noncatalytic functions that are sufficient for cell viability. Ser/Thr glycosylation requires residue D554 as a base to shuttle protons during the reaction, but this residue is not needed for HCF-1 proteolysis, which begins with glycosylation on a deprotonated glutamate side chain (Figure S1B) (24, 25, 50, 52). Mutating this residue therefore renders OGT inactive only for Ser/Thr glycosylation. OGT^D554N^ was therefore used to test if Ser/Thr glycosylation was required for survival. Because HCF-1 cleavage requires that the proteolytic repeat engage a network of five conserved asparagine residues in OGT’s tetratricopeptide repeat (TPR) domain (24), replacing these residues with Ala (OGT^5N5A^) completely blocks HCF-1 cleavage (24). Ser/Thr glycosylation is attenuated with the OGT^5N5A^ mutant (53) but is not abolished (Figure S1G). Therefore, even though OGT^5N5A^ is an imperfect separation-of-function variant, we thought it might allow us to determine if HCF-1 cleavage is necessary for cell survival. Below we refer to cell lines completely lacking OGT as *ΔOGT*, to those producing OGT^K842M^ as *OGT*^*ΔCat*^ for their lack of OGT catalytic activity, to those producing OGT^D554N^ as *OGT*^*ΔGlcNAc*^ for their lack of Ser/Thr glycosylation activity, and to those producing OGT^5N5A^ as *OGT*^*ΔClvg*^ for their lack of HCF-1 cleavage activity (Figure 2A).

**Figure 2:**
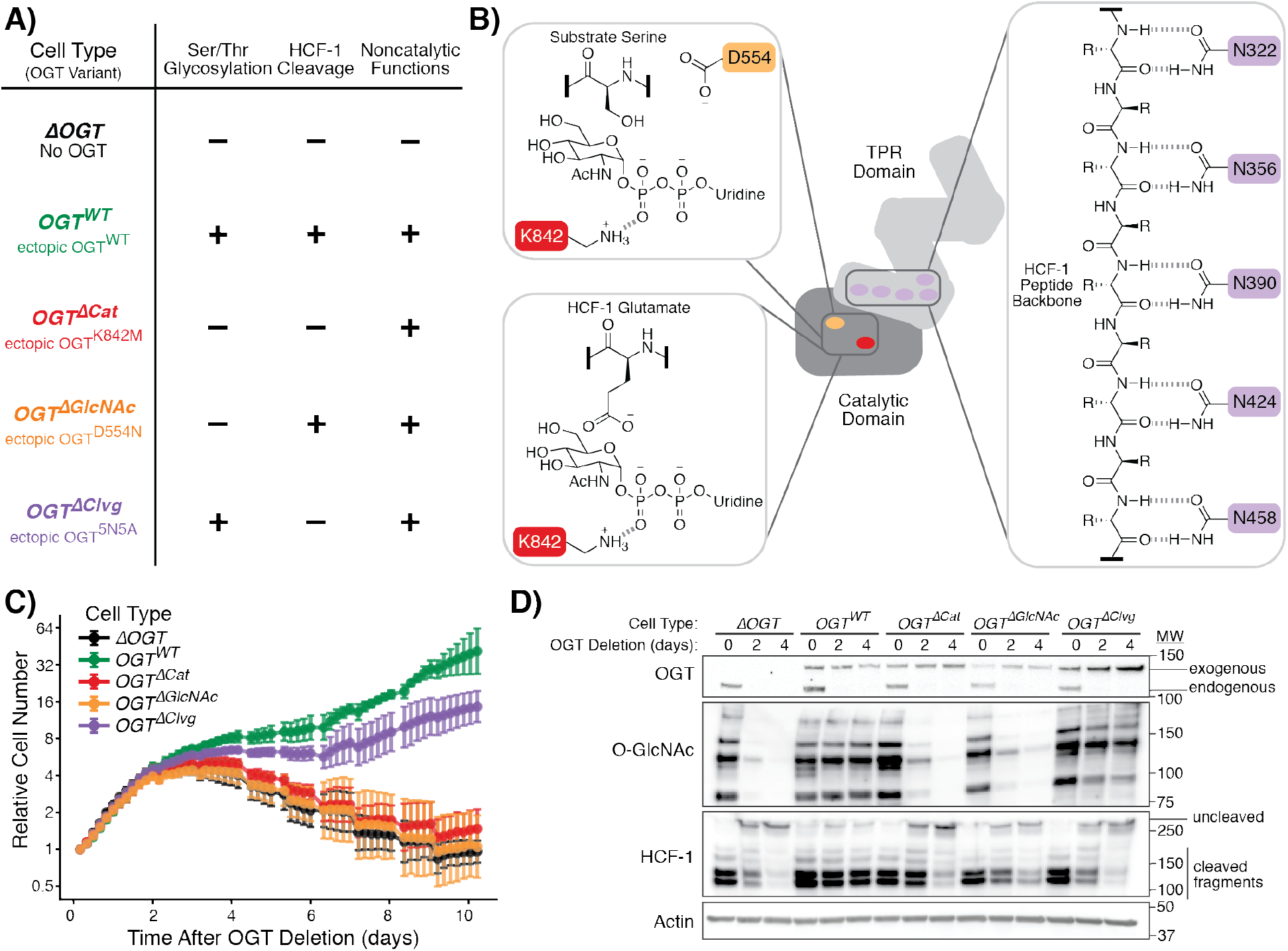
OGT’ s Only Essential Catalytic Function is Ser/ Thr Glycosylation. **A)** OGT variants with known *in vitro* activities can separate catalytic functions in living cells. Table shows cell type names and the associated OGT variant. “–” indicates absence of OGT activity, while “+” indicates presence of OGT activity. “5N5A” refers to mutation of 5 conserved TPR Asn residues to Ala. **B)** Mutation of key residues perturbs individual OGT activities. Active site residue K842 (red) is required for both Ser/Thr glycosylation (top left) and HCF-1 cleavage (bottom left). D554 (orange) mutation blocks Ser/Thr glycosylation but allows HCF-1 cleavage. Conserved TPR Asn residues (purple) bind substrates. Extensive binding is required for HCF-1 cleavage, but appreciable glycosylation is maintained upon mutation of five Asn residues to Ala (see Figure S1G). R= side chains. **C)** Cells without ability to add O-GlcNAc cease proliferation and die. Cells with *OGT* variants were imaged every 4 hours after inducing *Ogt* knockout at t=0. Relative cell number is based on normalized measurements of confluence from live cell imaging. Error bars are SD (n = 6 per condition). **D)** Immunoblot analysis confirms the expected loss of either one or both catalytic activities depending on the ectopically-produced OGT variant. See also Figure S1G.

We tested whether each of these OGT variants could complement loss of endogenous *Ogt*. Live cell microscopy showed that, like *ΔOGT* cells, *OGT*^*ΔCat*^ or *OGT*^*ΔGlcNAc*^ cells grew over the first two days, then underwent growth arrest and eventually died (Figure 2C). In contrast, *OGT*^*WT*^ and *OGT*^*ΔClvg*^ cells continued to proliferate. Immunoblot analysis showed that only *OGT*^*WT*^ and *OGT*^*ΔClvg*^ cells maintained protein O-GlcNAc levels (Figure 2D). The levels of ectopically-produced OGT increased over time in *OGT*^*ΔClvg*^ cells until they were noticeably higher than in other cell lines lacking *Ogt*. We concluded that *OGT*^*ΔClvg*^ cells with the highest ectopic OGT production outgrew *OGT*^*ΔClvg*^ cells with lower expression because higher OGT levels compensated for OGT^5N5A^’s attenuated Ser/Thr glycosylation activity. Even with this increased OGT production, HCF-1 cleavage was blocked in the *OGT*^*ΔClvg*^ cells (Figure 2D). The survival of these cells showed that HCF-1 cleavage is not required for cell viability. Because all cells that retained Ser/Thr glycosylation activity survived, whereas all cells without this activity died, Ser/Thr glycosylation is OGT’s only essential catalytic activity.

### Chemically-Induced Degradation Rapidly Separates OGT Functions

To study the direct cellular roles of the OGT variants, we sought to improve the speed of OGT depletion. Although genetic *Ogt* knockout is rapid, depletion of the endogenous OGT protein takes more than two days. By this time, cells completely lacking protein O-GlcNAc have undergone growth arrest and are in the process of dying. In addition, Cre recombinase induces DNA damage signaling unrelated to OGT activity (Figure S1F) (55). We were concerned that the cell death process and DNA damage signaling would obscure direct cellular roles of OGT’s catalytic activities. We speculated that if we could rapidly deplete wild-type OGT in the presence of a variant that retained only some activities, we might be able to deconvolute initial responses due to loss of an activity from other effects. Moreover, a rapid depletion system to synchronously alter cells’ OGT activities might allow study of OGT variants that do not contain the full repertoire of functions required for viability.

To deplete wild-type OGT rapidly, we introduced an FKBP12^F36V^-OGT fusion into the inducible *Ogt* knockout MEFs (Figure S2A) to enable chemically-induced protein degradation (56). FKBP12^F36V^ provides a binding site for one half of the heterobifunctional degrader molecule dTAG-13 (56). The other half of dTAG-13 recruits the E3 ligase cereblon, leading to ubiquitination and proteasomal degradation of FKBP12^F36V^-OGT upon treatment (Figure 3A). After knocking out endogenous *Ogt*, we isolated multiple MEF clones expressing only *FKBP12*^*F36V*^-*OGT* (Figure 3B). These clones were viable several weeks after endogenous *Ogt* removal, demonstrating that the FKPB12^F36V^ tag did not interfere with essential OGT functions. Treatment of *FKBP12*^*F36V*^-*OGT* cells with dTAG-13 depleted OGT within one hour with depletion sustained for 24 hours (Figure 3C, inset). We observed dose-dependent growth inhibition with maximal inhibition by 500 nM dTAG-13 (Figure 3C), a concentration used for selective degradation of tagged proteins in several prior studies (56–58). Growth inhibition implied that OGT levels were sufficiently reduced to block OGT’s growth-promoting cellular functions, providing a background in which we could assess how OGT’s individual activities control cell physiology.

**Figure 3:**
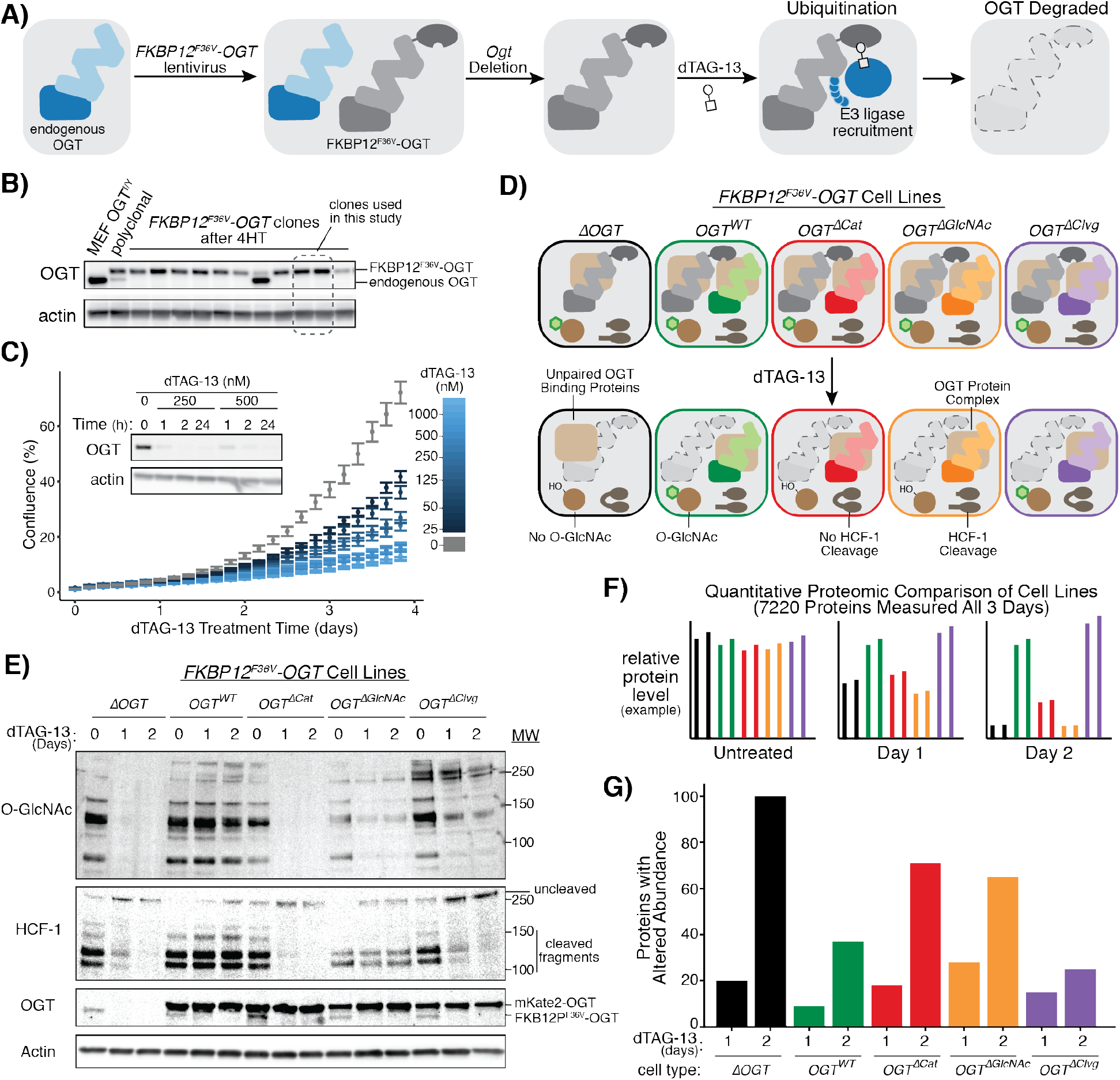
Rapid OGT Degradation Reveals Protein Level Changes Due to Loss of Specified Activities. **A**) The dTAG system enables rapid, inducible OGT depletion. A copy of OGT tagged with FKBP12^F36V^ is used to replace endogenous OGT in MEFs. Treatment with heterobifunctional molecule dTAG-13 recruits endogenous E3 ligase cereblon to ubiquitinate FKBP12^F36V^-OGT, leading to degradation. **B)** Clonal MEFs containing FKBP12^F36V^-OGT were isolated at 2 weeks after *Ogt* knockout. Two clones lacking endogenous OGT were selected for further study. **C)** FKBP12^F36V^-OGT is rapidly degraded, leading to growth arrest. dTAG-13 treatment leads to dose-dependent growth arrest as measured by live cell imaging (n=6, n=3 per clone). Error bars are standard error of the mean. Sub-micromolar doses of dTAG-13 lead to rapid, durable degradation of FKBP12^F36V^-OGT by western blot analysis (inset). **D)** Combining FKBP12^F36V^-OGT with separation-of-function OGT variants allows synchronous separation of OGT activities. dTAG-13 treatment “unmasks” separation of function by rapidly removing wild-type OGT, allowing analysis of cells as specified OGT functions are lost. **E)** Western blot analysis demonstrating separation of function within one day for *FKBP12*^*F36V*^-*OGT* cells. mKate2-OGT bands show levels of OGT variants specifically added to each cell type. **F)** Quantitative mass spectrometry profiles how altered OGT biochemistry changes protein levels proteome-wide. Isobaric peptide labeling and LC/MS^3^-based protein quantitation enabled measurement of 7220 proteins’ levels across samples treated with dTAG-13 for 0, 1, or 2 days. Graphs represent hypothetical data. **G)** More proteins changed levels with loss of OGT protein than with loss of its catalytic functions. Graph shows number of hits for each cell type after one or two days dTAG-13 treatment relative to untreated control (hits: >1.5 fold change in level and a p-value < 0.01 in a t-test). See also Figure S2.

We next introduced the separation-of-function OGT variants into clonal FKBP12^F36V^-OGT cell lines (Figure 3D). All cells had wild-type OGT activities before treatment, but at 24 and 48 hours after dTAG-13 treatment, we only detected the OGT activities expected for each variant (Figure 3E). Consistent with the OGT^5N5A^ mutant being partially impaired in Ser/Thr glycosylation, protein O-GlcNAc levels were moderately reduced in the *OGT*^*ΔClvg*^ cells after dTAG-13 treatment.

We measured relative protein abundance in the different cell lines to query changes in cell physiology when specified OGT activities were removed (Figure 3F, Figure S2B). Two biological replicates of each of our five FKBP12^F36V^-OGT cell types (*ΔOGT*, *OGT*^*WT*^, *OGT*^*ΔCat*^, *OGT*^*ΔGlcNAc*^, and *OGT*^*ΔClvg*^) were treated with dTAG-13 for 1 and 2 days; an untreated control (Day 0) was also collected, and samples were labeled with isobaric tags to enable relative protein quantitation (59–61). LC/MS quantified 7220 distinct proteins in all samples at all three timepoints (Dataset S1, Figure S2B). For all cell lines, dTAG-13 treatment altered levels of more proteins at Day 2 than Day 1 (as measured by proteins changing >1.5 fold with a p-value of <0.01 in a two-sample t-test, Figure 3G, Dataset S2). At Day 2, the fewest changes were observed for *OGT*^*WT*^ and *OGT*^*ΔClvg*^ cells (37 and 25 proteins with altered abundance, respectively) and the most were observed for *ΔOGT* cells (100 proteins with altered abundance). Cells expressing an OGT variant incapable of Ser/Thr glycosylation (*OGT*^*ΔGlcNAc*^ and *OGT*^*ΔCat*^) had an intermediate number of altered proteins (65 and 71). These results showed that Ser/Thr glycosylation plays a larger role in regulating protein levels than HCF-1 cleavage. Because there were more changes in *ΔOGT* than in the *OGT*^*ΔCat*^ cells, the results also suggested that OGT may have physiologically important noncatalytic functions.

### Simultaneous Comparison of Multiple Cell Lines Quantifies Effects of Individual OGT Activities

To use the proteomics data to identify proteins affected by individual OGT activities, we needed an analytical method to link changes in protein levels to the activities driving the changes. We reasoned that if a protein’s abundance depended only on a single OGT activity, then the levels of that protein should differ based on whether a given cell line retained the activity after dTAG-13 treatment (Figure 4A). For example, levels of proteins controlled by Ser/Thr glycosylation or HCF-1 cleavage should differ between the two cell lines that retained that particular catalytic activity and the three cell lines without it. If a protein’s abundance primarily depended on a noncatalytic function of OGT, levels should be similar for the four cell lines producing a copy of OGT but different in *ΔOGT* cells. The log_2_ fold change in abundance of a protein upon losing an activity is the “effect size” of that OGT activity on that protein. For example, increased abundance of a protein when Ser/Thr glycosylation is blocked would lead to a positive Ser/Thr glycosylation effect size, whereas decreased abundance would produce a negative Ser/Thr glycosylation effect size. We hypothesized that we could determine which proteins depended on which OGT activities by modeling protein levels as a linear combination of activity-specific effect sizes. Therefore, for each protein in the proteome, we used linear regression to calculate *β*_*GlcNAc*_, *β*_*Clvg*_, and *β*_*Ncat*_, which are the activity-specific effect sizes that, when added together, describe how levels of that protein change when a specific set of OGT activities are removed (Figure 4B, Dataset S3). By allowing us to use data from all five cell lines simultaneously, this approach filtered out cell line-specific effects – for example, protein level changes due to reduced glycosylation in the *OGT*^*ΔClvg*^ cells – because these would result in noisy effect sizes that did not meet p-value cutoffs for statistical significance.

**Figure 4:**
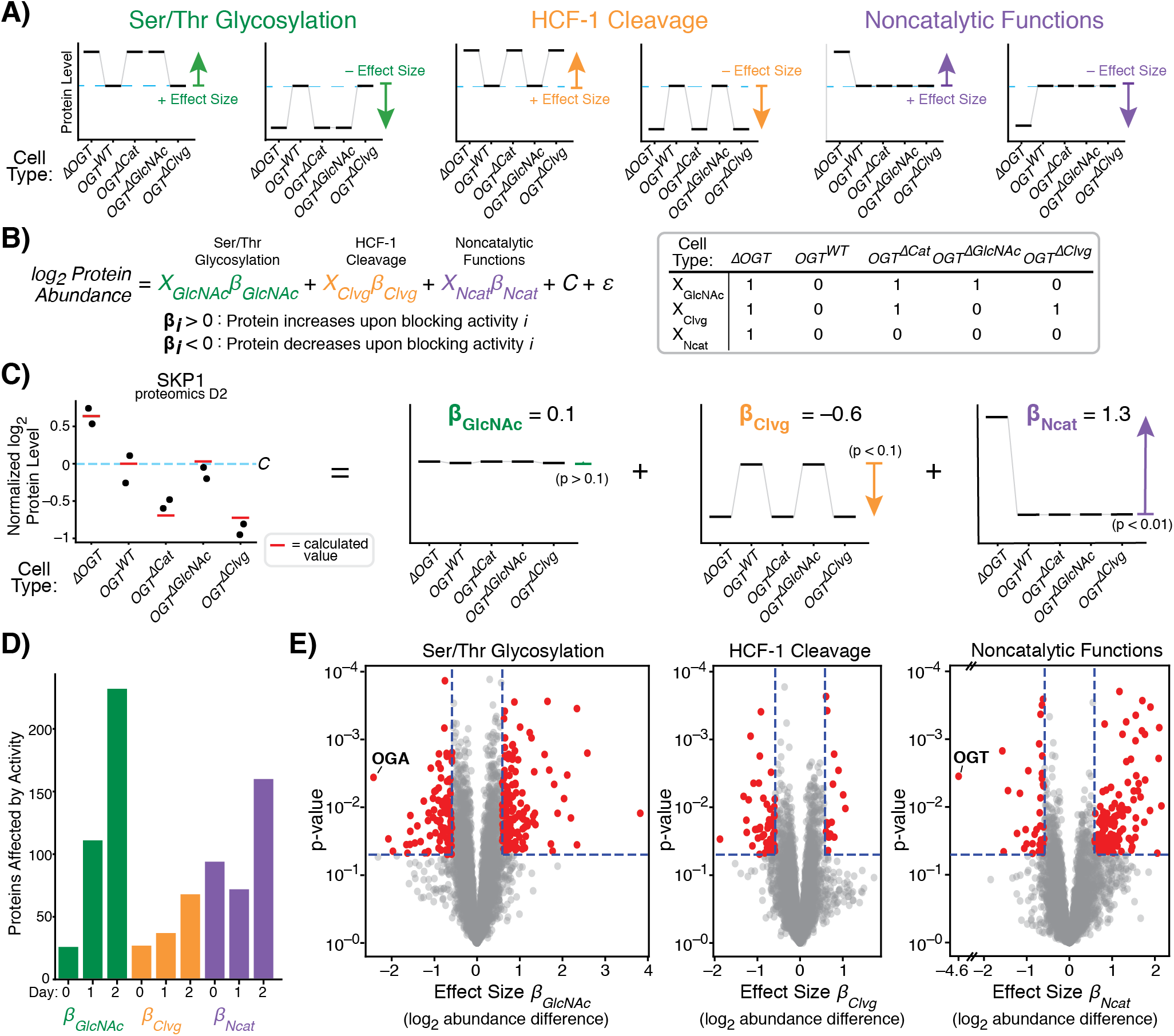
Regression Measures How Single OGT Activities Alter Levels of Each Protein. **A)** A protein whose levels are controlled by one OGT activity will show a characteristic patterns amongst the 5 cell lines. Each graph represents a hypothetical protein controlled by a single activity; examples are shown for proteins that increase or decrease in response to inhibiting each activity. Protein level in *OGT*^*WT*^ cells shown by blue dashed line. **B)** Linear regression quantifies degree to which protein levels match the patterns corresponding to a given OGT activity. *β*_*GlcNAc*_, *β*_*Clvg*_, and *β*_*Ncat*_ are log_2_ fold-change upon inhibiting an activity, and *X*_*GlcNAc*_, *X*_*Clvg*_, and *X*_*Ncat*_ indicate which cell lines lose a given OGT activity upon dTAG-13 treatment. *C* is average level in *OGT*^*WT*^ cells and *ε* is random variation unrelated to OGT activity. **C)** Protein SKP1 is shown as a regression example; dots are measured abundance, red lines are calculated from linear regression, and individual effects are shown at right. p-values shown for each effect. Level in *OGT^WT^* cells shown by blue dashed line. **D)** Number of hits for each day for each OGT activity. (Hits: p < 0.05 and effect size > log_2_ (1.5)) **E)** Volcano plot showing *β*_*GlcNAc*_, *β*_*Clvg*_, and *β*_*Ncat*_ from 2 days of dTAG-13 treatment. OGT and OGA are highlighted as expected outliers. See also Figures S3 and S4.

We observed that levels of some proteins depended on a single OGT activity (Figure S3A) whereas others were best represented as a combination of OGT activities. For example, coactosin-like 1 (COTL1), an actin-binding protein, was less abundant in cells lacking HCF-1 cleavage activity (Figure S3A, left), whereas caldesmon 1 (CALD1), a protein involved in calcium-dependent actomyosin regulation, was more abundant in cells without a copy of OGT (Figure S3A, middle). The protein chaperone α-crystallin B (CRYAB), known to be O-GlcNAcylated and previously shown to be upregulated in response to OGT knockout (40, 62), increased in cells lacking Ser/Thr glycosylation activity (Figure S3A, right). Therefore, we can attribute the increase in CRYAB abundance upon OGT knockout to loss of O-GlcNAc. In contrast to these proteins, which were dependent on a single OGT activity, SKP1, a protein in the SCF ubiquitin ligase complex, was sensitive to both loss of OGT, which led to increased abundance (*β*_*NCat*_ = 1.3), and to loss of HCF-1 cleavage, which led to decreased abundance (*β*_*Clvg*_ = −0.6), but not to loss of Ser/Thr glycosylation (Figure 4C). By using this linear regression analysis, we could therefore demonstrate that HCF-1 cleavage and noncatalytic OGT functions both regulate SKP1 levels, with a larger role for the noncatalytic functions.

Several pieces of data showed that effect sizes generated across all three time points (Figures S3B–C, Dataset S4) accurately reflected how levels of each protein responded to loss of individual OGT activities. Proteome-wide correlation analysis showed that effect sizes for loss of each OGT activity corresponded to the differences in protein abundance between any two sets of samples that differed by that OGT activity, both between different cell lines (Figure S4A, Dataset S5) and between untreated and treated samples of the same cell line (Figure S4B). Expected outliers in the data further validated the regression analysis (Figure 4E). For example, OGA abundance is known to decrease with low protein O-GlcNAc levels (40, 44), and we found that OGA had the most negative *β*_*GlcNAc*_ value after two days of dTAG-13 treatment. Furthermore, OGT had a negative *β*_*Ncat*_ value that corresponded to a striking 24-fold reduction in protein abundance; this result was reassuring because noncatalytic function effect sizes are defined by the presence or absence of OGT. We concluded that the regression analysis was an appropriate method to identify the proteins most affected by individual OGT activities.

We made two observations about the numbers of affected proteins and their changes over time (Figure 4D). First, on Day 0, few proteins were affected by Ser/Thr glycosylation or HCF-1 cleavage. We inferred that the presence of catalytically-active FKBP12^F36V^-OGT obscures differences in catalytic activity from other variants. In contrast, a substantial number of proteins were affected by noncatalytic functions at Day 0. Noncatalytic roles are likely to involve stoichiometric binding interactions, and a second copy of OGT at Day 0 would provide more protein to fulfill these roles. Second, after dTAG-13 treatment, loss of HCF-1 cleavage affected levels of a relatively small number of proteins compared to loss of OGT’s other activities, implying a narrower role for this activity in regulating cell physiology. Loss of Ser/Thr glycosylation and noncatalytic functions each altered levels of more than 150 proteins by Day 2, demonstrating a much larger impact of these activities.

### Noncatalytic Functions and Ser/ Thr Glycosylation Control Levels of Different Proteins

Proteome-wide analysis of the effect sizes indicated that each OGT activity altered the levels of different sets of proteins (Figure S5A) and revealed that the majority of hit proteins had large effect sizes for only one OGT activity (Figure 5A). Supporting the idea that different proteins are affected by different OGT activities, we find enrichment of previously-identified O-GlcNAc proteins among proteins with significant Day 2 effect sizes for Ser/Thr glycosylation, and enrichment of OGT interactors in proteins with significant effect sizes for noncatalytic functions (Table S1) (33, 63). To examine the processes altered by individual OGT activities, we performed gene set enrichment analysis (GSEA) based on the effect sizes and their degree of statistical significance (Figures 5B, S5D, Dataset S7) (64–66) to identify pathways affected by the different activities. Each OGT activity primarily affected different pathways from the others, although several pathways were affected by multiple OGT activities. Notably, OGT’s noncatalytic function(s) were associated with oxidative phosphorylation and actin cytoskeleton components (Figure 5B, S5D–E), while OGT’s catalytic activities were associated with other areas of metabolism and numerous aspects of gene expression. Overall, the analysis showed that OGT’s different biochemical activities – Ser/Thr glycosylation, HCF-1 cleavage, and noncatalytic functions – control different aspects of cell physiology.

**Figure 5:**
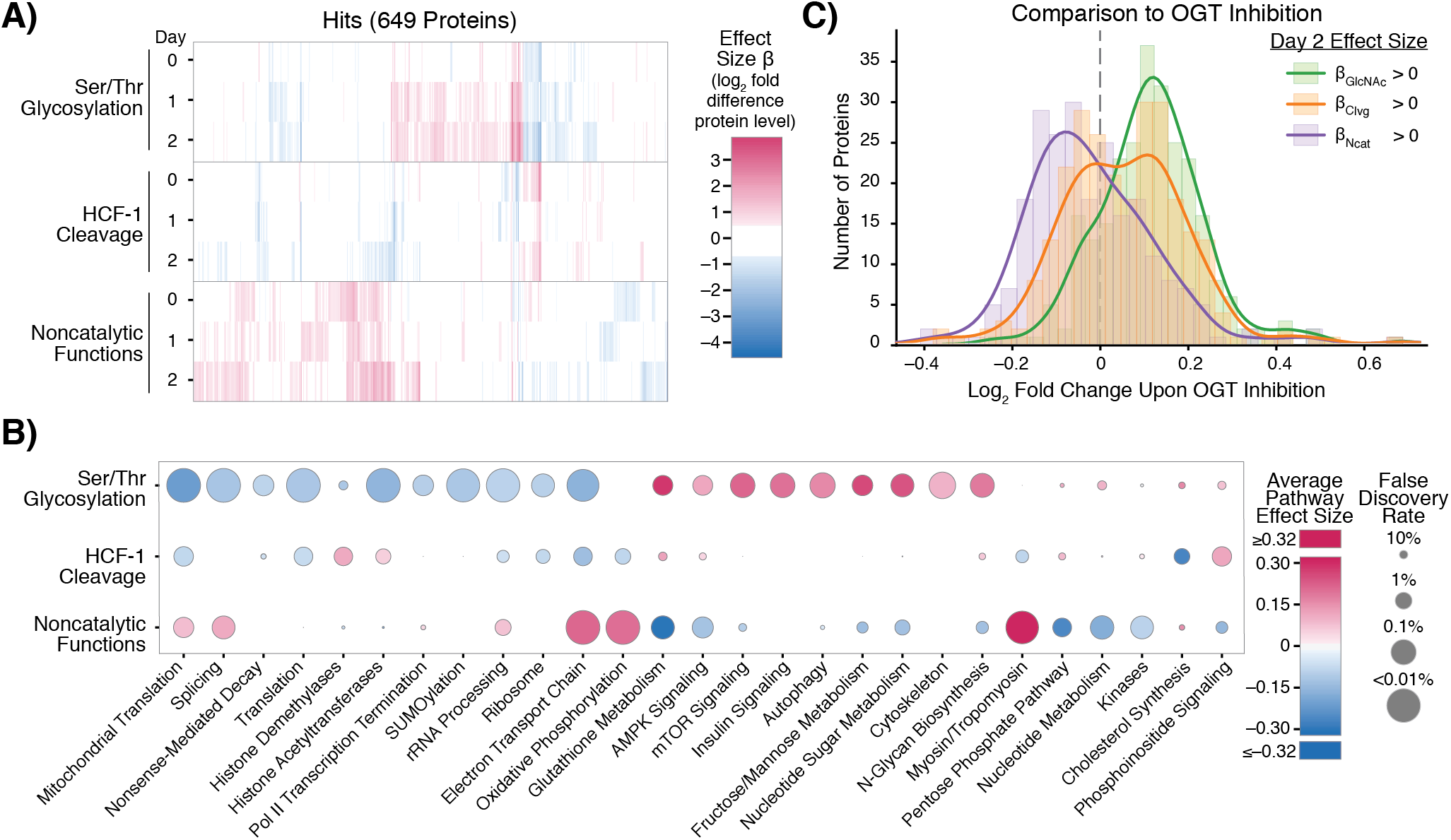
Ser/ Thr Glycosylation and Noncatalytic Functions Control Distinct Cellular Processes. **A)** Heatmap of effect sizes for all 649 protein hits for any OGT activity at any length of dTAG-13 treatment. OGT activity and length of dTAG-13 treatment shown on Y-axis. Proteins arrayed on X-axis. **B)** OGT activities control different pathways. Average effect sizes of all proteins (up to 25% change, effect size 0.32) in a given pathway and false discovery rate are based on each OGT activity after two days dTAG-13 treatment. Size of points corresponds to false discovery rate from GSEA.See also Figure S5D. **C)** Comparison of OGT inhibition and changes due to individual OGT activities. Histograms show log_2_ fold change in protein abundance upon OGT inhibition (Martin et al. 2018) for the set of proteins with positive Day 2 β_GlcNAc_ (n = 263), β_Clvg_ (n = 264), and β_Ncat_ (n = 235) values. Lines show smoothed distribution for each set of proteins (gaussian kernel with bandwidth of 0.05). Only proteins significant in regression analysis (p<0.01 for any effect size, 513 proteins) were analyzed. Proteins with positive effect sizes upon blocking any individual OGT activity had significantly more extreme changes in abundance upon OGT inhibition than expected by chance based on a Wilcoxon rank sum test (p-value β_GlcNAc_ < 10^−26^, β_Clvg_ < 0.01, β_Ncat_ < 10^−10^). See also Figures S5B and S6

We also compared the effects of altering individual OGT activities to the effects of inhibiting OGT with a small molecule. Using a previously-reported data set for OGT inhibition, we compared the changes in protein abundance observed after treating human HEK 293T cells with an OGT active site inhibitor to the effect sizes obtained from our genetic perturbations in MEFs (Figure S6, Dataset S6) (67). Overall, we found that protein levels in HEK 293T cells changed with OGT inhibition in the same direction as their mouse orthologues did when OGT’s catalytic activities were removed (Figures 5C, S5B). The correspondence between protein abundance changes upon OGT active site inhibition and effect sizes for loss of OGT’s catalytic activities provided independent evidence that these effect sizes are useful metrics for quantifying the contributions of OGT’s catalytic activities to regulating protein abundance. Moreover, because inhibiting OGT in HEK 293T cells and removing OGT’s catalytic activities in MEFs affected similar biological pathways (Figure S5C–D, Dataset S7), we have inferred that OGT’s catalytic activities play consistent roles in controlling cell physiology in different mammals.

Unexpectedly, we also observed a relationship between OGT inhibition and OGT’s noncatalytic functions. Inhibition altered protein levels in the *opposite* direction from removing OGT’s noncatalytic functions, a phenomenon observed at both the protein and biological pathway levels (Figures 5C, S5B–D). It is well-known that OGT levels rapidly increase upon loss of protein O-GlcNAcylation (44, 48, 67, 68), with the rapid increases driven by splicing and transcriptional controls that we removed in the MEF system. These mechanisms were thought to serve as a means to restore normal O-GlcNAc levels. However, we interpret the inverse correlation between protein-level changes after small molecule OGT inhibition and effect sizes for removing OGT’s noncatalytic functions as evidence that increased OGT abundance is important independent of altering OGT catalytic activity. Therefore, OGT’s noncatalytic functions contribute to the normal cellular response to reduced O-GlcNAcylation.

### Noncatalytic Functions are Required for Cell Proliferation

Given that our analysis strongly implicated noncatalytic functions of OGT in controlling cell physiology, we wondered if these functions were important for cell growth. To test if OGT’s noncatalytic activities contribute to cell proliferation, we tracked growth of our five FKBP12^F36V^-OGT cell lines in the presence of enough dTAG-13 to suppress proliferation of *ΔOGT* cells (Figure 6A). As expected, the *OGT*^*WT*^ and *OGT*^*ΔClvg*^ cells, which can carry out Ser/Thr glycosylation, grew in the presence of the degrader compound; however, the *OGT*^*ΔGlcNAc*^ and *OGT*^*ΔCat*^ cells, which are nonviable in the Cre-based deletion system, also grew (Figure 6A–B). Because the amounts of catalytically-active FKBP12^F36V^-OGT remaining after dTAG-13 treatment are too low to permit cell proliferation, growth of the *OGT*^*ΔCat*^ cells requires noncatalytic roles of OGT (Figure 6C).

**Figure 6:**
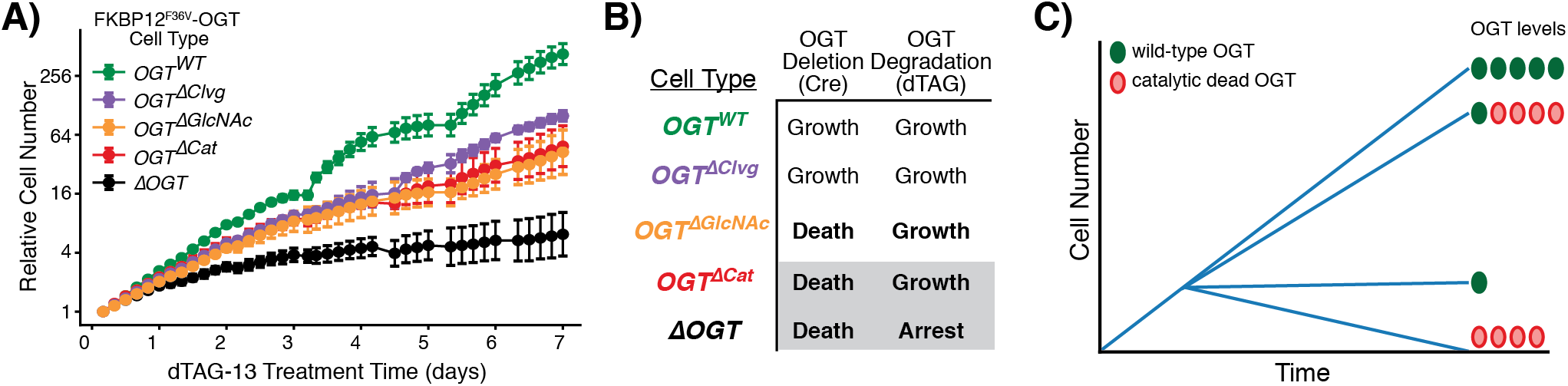
OGT’ s Noncatalytic Roles are Required for Cell Proliferation. **A)** Cell growth over time of all FKBP12^F36V^-OGT cell lines, as measured by live cell imaging. dTAG-13 treatment began at t=0. Relative cell number is based on normalized measurements of confluence from live cell imaging. Error bars are SD (n = 6 per condition). **B)** Growth outcomes for the Cre-based OGT deletion (Figure 2) and dTAG-based OGT degradation systems (Figure 6A). The degron-tagged *ΔOGT* cell line is viable because a very small amount of OGT remains after dTAG-13 treatment. **C)** Schematic illustrating that OGT integrates catalytic and non-catalytic functions to promote cell growth.

## Discussion

This study identified the OGT functions that are essential for growth of mammalian cells. First, by replacing OGT with variants impaired in selected activities, we determined that protein O-GlcNAcylation is required for cell survival. HCF-1 cleavage is not. Next, by using inducible protein degradation and a broadly applicable proteomic analysis method to deconvolute the roles of variants that are deficient in one or more OGT activities, we demonstrated that OGT has noncatalytic roles that regulate protein levels independent of catalysis. We also showed that these noncatalytic functions are sufficient to drive growth when there is insufficient wild-type OGT to support cell proliferation. Taken together, our results show that OGT has two essential functions, Ser/Thr glycosylation and a noncatalytic role.

OGT’s catalytic and noncatalytic functions evidently cooperate to control cell proliferation. If no OGT is present, or if OGT is present but cannot catalyze any Ser/Thr glycosylation, cells die. If wild-type OGT is present, but at extremely low levels, cells are viable but remain in a growth-arrested state. Adding back catalytically-inactive OGT rescues cell proliferation even though protein O-GlcNAc levels remain low. Therefore, providing sufficient OGT to fulfill its stoichiometric noncatalytic roles promotes cell proliferation even if only a small amount of O-GlcNAc is present. It remains to be established how OGT’s noncatalytic functions promote cell proliferation, but possibilities include a role in binding other proteins to stabilize them, to alter their localization, or to scaffold the formation of larger complexes.

The importance of OGT’s noncatalytic functions for cell growth helps explain a previously observed discrepancy between genetic OGT knockout and OGT inhibition. Cells die upon knockout (40) but not when OGT is inhibited. Death occurs in the former case because both O-GlcNAc signaling and OGT’s noncatalytic functions are removed. With inhibition, low levels of Ser/Thr glycosylation persist. The reduced protein O-GlcNAc levels trigger a rapid increase in OGT abundance through altered splicing and transcription (44, 48, 67). When the increase in OGT abundance is sufficient to overcome inhibition, O-GlcNAc levels recover. Whether or not they recover, signaling through OGT’s noncatalytic functions increases because there is more OGT protein available. Noncatalytic functions drive cell proliferation even when O-GlcNAc levels remain very low.

OGT’s noncatalytic functions may be important for OGT’s role as a nutrient sensor. Low nutrients lead to reduced cytosolic UDP-GlcNAc levels (8, 22), leading to a transient reduction in O-GlcNAc levels that is followed by rapid upregulation of OGT (68). If protein O-GlcNAc levels remain low even though OGT abundance is high, noncatalytic signaling may become especially important. We suggest that OGT uses both its catalytic and noncatalytic functions to respond to environmental changes, which may allow for varied responses to different cellular cues.

By quantifying how much each OGT activity impacts the abundance of each protein in the cell, we have identified the biological pathways most affected by each OGT activity. Consistent with known roles for O-GlcNAc, we found that Ser/Thr glycosylation broadly regulates metabolism and gene expression, with enriched pathways including splicing (44), translation (69–71), and autophagy (72–74). Among the pathways linked to OGT’s noncatalytic activities are oxidative phosphorylation, the electron transport chain, and components of the actin cytoskeleton, consistent with prior data suggesting OGT binds to the actin-regulating E-cadherin/beta-catenin complex (39). We also note that a previous report showed that proteins involved in oxidative phosphorylation and electron transport were downregulated upon OGT overexpression (75), consistent with our finding that blocking OGT’s noncatalytic functions leads to an increase in the levels of these proteins. These previous studies attributed altered abundance of proteins involved in respiration to increased O-GlcNAc, but our results suggest that OGT’s noncatalytic functions also contribute substantially to altering levels of these proteins.

Our data show that OGT’s glycosyltransferase and noncatalytic OGT functions can cooperate within the same cellular pathway to alter cell physiology. When nutrients are low or OGT is inhibited, signaling through Ser/Thr glycosylation is reduced, but elevated OGT levels lead to increased signaling through OGT’s noncatalytic functions. Our effect sizes measure the response of protein levels to *loss* of an OGT activity so an increase in noncatalytic signaling would lead to protein-level changes in the opposite direction from the noncatalytic effect sizes. As a result, effect sizes with opposite signs for Ser/Thr glycosylation and noncatalytic functions indicate protein level changes in the same direction. Our analytical method identifies pathways dominated by one activity as well as pathways where both catalytic and noncatalytic OGT functions substantially contribute to regulation (Figure 5). Going forward, it will be critical to determine whether effects of genetic knockdown are due to loss of O-GlcNAc, loss of OGT’s noncatalytic functions, or both.

HCF-1 cleavage affected a more limited number of cellular processes than OGT’s other activities, but it remains a central player in OGT biology. We found that the cleavage activity affects histone modification pathways, which include known interactors of HCF-1 (76, 77), as well as lipid metabolism and peroxisome-related processes (Figure 5C, S5D, Dataset S7), which have been linked to HCF-1 based on hepatocyte-specific HCF-1 knockout mice (78). Because HCF-1 is one of OGT’s most common cellular binding partners (26, 29, 32, 33, 77) and is O-GlcNAcylated in addition to being proteolyzed, it integrates all three of OGT’s biochemical activities. HCF-1 regulates transcription through participation in several chromatin-modifying complexes (33, 77, 79), and recent evidence suggests it is directly involved in altering gene expression in response to nutrient cues, including cues that result in altered O-GlcNAc (32, 33, 80). Moreover, both O-GlcNAc modification and cleavage of HCF-1 have been shown to change its binding partners and consequently alter downstream gene expression (32, 81). Effects of HCF-1 cleavage were previously investigated by altering HCF-1 structure. The ability to disrupt cleavage by varying OGT rather than by altering HCF-1 may have some advantages for understanding the role of HCF-1 in gene expression and the interplay of cleavage with HCF-1 O-GlcNAcylation and OGT binding.

We expect the genetic systems and approaches described here will be useful for addressing longstanding questions about OGT. One important question is how O-GlcNAcylation supports cell viability. It is possible that a small number of OGT substrates require O-GlcNAc modification to carry out essential functions. If so, these might be found by identifying the proteins that are still modified by OGT variants that have lost the ability to glycosylate most substrates. Similarly, the ability to introduce OGT variants into cells and either delete or deplete wild-type OGT will help elucidate the binding partners through which OGT engages in noncatalytic signaling. These binding partners likely include other proteins, but may also include nucleic acids or other molecules. OGT’s TPR domain has been implicated in binding interactions (3, 34), and the ability to modify this domain to disrupt binding interactions should be helpful in dissecting OGT function. The approaches and analytical strategy we have developed to quantify effects of OGT’s activities on protein levels can also be adapted to understand other multifunctional proteins. Moreover, although we used protein abundance to quantify effect sizes in the studies here, other quantitative phenotypes such as transcriptomic measurements can also be used to link particular functions to pathways (82). Last, by extending these approaches to alternative cell types, we anticipate that we may be able to determine which effects of OGT’s activities are general and which are only relevant in the signaling context of a particular cell type.

Finally, we note that the results here have implications relevant to OGT-targeting therapeutics. OGT has been identified as a potential target in multiple diseases including cancer (14–17, 47, 49) and metabolic disease (8, 18–21). Many of the studies implicating OGT in these diseases used genetic perturbations that blocked catalytic and noncatalytic signaling simultaneously. However, OGT deletion and depletion phenotypes are different from phenotypes due to OGT active site inhibition. Anti-proliferative effects are more likely to occur from removing OGT entirely than from blocking its active site because, as we have shown, cells can proliferate with very low catalytic activity if there is sufficient OGT protein to fulfill noncatalytic roles. Therefore, targeting OGT for cancer therapy may require either a targeted protein degradation approach (83) that disrupts both catalytic and noncatalytic OGT activities or a strategy that exploits synthetic lethality between OGT inhibition and another pathway (84). By contrast, if metabolic syndrome indeed results from excess Ser/Thr glycosylation, it may be possible to use OGT inhibitors chronically to reduce protein O-GlcNAc levels; however, inhibitor treatment may lead to chronic elevation of OGT levels, and our findings show that a persistent increase in OGT may affect cell physiology independent of catalysis.

## Materials and Methods

### Plasmids and Cloning

Plasmids for protein expression and lentivirus production were generated as described in SI Text. Mutations were introduced by Q5 mutagenesis or Gibson assembly.

### Cells and Lentivirus

MEF cell line was maintained as previously described (40). Lentivirus was produced in HEK293T cells and infections of MEF cells were carried out as described in SI Text. Knockout of OGT was performed as previously described (40). Degradation of FKBP12^F36V^-OGT was carried out by changing to media containing 500 nM dTAG-13. Cell lysis and western blot analysis were carried out as described in SI Text.

### Growth Assays

All growth assays were carried out in 96 well plates using the Essen Incucyte Zoom platform as described in SI Text. Relative cell number was based upon confluence normalized to initial confluence. To enable long-term tracking of growth, cells that reached 85% confluence were split 1:3 and re-plated for continued growth, with relative cell number reflecting continued growth from point of splitting.

### Quantitative Proteomics

Samples were prepared, labeled with isobaric TMT labels, fractionated by HPLC, and analyzed by LC-MS^3^ as previously described (59–61), with further details in SI Text. A normalization sample of peptides from all other samples was generated by add 3 μL each of all 30 samples to the same tube before TMT labeling. LC-MS data was analyzed as described in SI Text to yield relative protein abundance. For analysis across days, each protein’s abundance was divided by protein abundance in the normalization sample for that day. *Linear Regression Analysis* Linear regression as described in Figure 4B was used to analyze each protein. Samples were randomly shuffled 10^6^ times to generate background distributions to determine p-value for each regression coefficient (effect size). For linear regression across multiple days, ten coefficients (3 days of 3 effect sizes plus a constant representing baseline abundance) were included with 30 samples, with the same procedure used to generate p-values. Further details are available in SI Text.

### Pathway Analysis

Gene Set Enrichment analysis was carried out using either effect-size or log_10_ of p-value (negative log_10_ if effect size was positive). Pathways were considered significant if they met a 10% FDR cutoff in both analyses and a 1% cutoff in either analysis. Further details are provided in SI Text.

### Comparison to OGT Inhibitor

Data from Martin, Tan et al. (67) as presented in SI of that publication was aligned on the basis of gene orthology and analyzed as described in text. Further details of analysis available in SI Text.

### Materials Availability

Request for materials or data should be directed to. All plasmids generated in this study are available through Addgene as described in Table S3, and code for analysis is available on Github at https://github.com/SuzanneWalkerLab/NoncatalyticOGT/.

## Acknowledgements

We thank Prof. Connie Cepko for advice on lentiviral systems and the Immunology Flow Cytometry Core Facility at Harvard Medical School for assistance with cell sorting. We thank Prof. Steven P. Gygi for providing access to facilities for mass spectrometry experiments. This work was supported by the NIH (GM094263 to S.W.; GM132129 to J.P.; K12HL141952 & CA230978 to N.E.Z., and T32 GM095450 to Z.G.L., S.J.P, and G.Q.F.) and the Katherine L. and Steven C. Pinard Research Fund (N.S.G.), with additional support in the form of pre- and postdoctoral fellowships (National Science Foundation grant DGE1745303 to S.C.P.; F32 GM129889 to C.M.J.; 5F31GM116451 to M.S.; American Cancer Society Postdoctoral Fellowship PF-17-010-01-CDD to B.N.). Aid and equipment for live cell imaging were provided by Dr. Clarence Yapp and the Laboratory for Systems Pharmacology at Harvard Medical School, which is supported in part by NIH grants P50GM107618 and U54CA225088, DARPA grant W911NF-19-2-0017, and the Ludwig Center at Harvard.

## Author Contributions

Conceptualization, Z.G.L. and S.W.; Methodology, Z.G.L., J.A.P.; Formal Analysis, Z.G.L. and M.S.; Investigation, Z.G.L., S.C.P., C.M.J., G.Q.F.; Writing – Original Draft, Z.G.L., S.C.P., S.W.; Resources, B.N., N.E.Z., N.S.G., J.A.P.; Funding Acquisition, S.W.; Supervision, S.W.

## References

1. M. Jöud, M. Möller, M. L. Olsson, Identification of human glycosyltransferase genes expressed in erythroid cells predicts potential carbohydrate blood group loci. Sci. Rep. 8, 6040 (2018).

2. K. J. Karczewski, et al., The mutational constraint spectrum quantified from variation in 141,456 humans. Nature 581, 434–443 (2020).

3. Z. G. Levine, S. Walker, The Biochemistry of O-GlcNAc Transferase: Which Functions Make It Essential in Mammalian Cells? Annu. Rev. Biochem. 85, 631–657 (2016).

4. N. O’Donnell, N. E. Zachara, G. W. Hart, J. D. Marth, Ogt-dependent X-chromosome-linked protein glycosylation is a requisite modification in somatic cell function and embryo viability. Mol Cell Biol 24, 1680–1690 (2004).

5. R. Shafi, et al., The O-GlcNAc transferase gene resides on the X chromosome and is essential for embryonic stem cell viability and mouse ontogeny. Proc. Natl. Acad. Sci. 97, 5735–5739 (2000).

6. N. Zachara, Y. Akimoto, G. W. Hart, “The O-GlcNAc Modification” in Essentials of Glycobiology, 3rd Ed., A. Varki, et al., Eds. (Cold Spring Harbor Laboratory Press, 2015) (February 6, 2020).

7. R. Trapannone, D. Mariappa, A. T. Ferenbach, D. M. F. van Aalten, Nucleocytoplasmic human O-GlcNAc transferase is sufficient for O-GlcNAcylation of mitochondrial proteins. Biochem. J. 473, 1693–1702 (2016).

8. X. Yang, K. Qian, Protein O-GlcNAcylation: emerging mechanisms and functions. Nat. Rev. Mol. Cell Biol. 18, 452–465 (2017).

9. D. L. Dong, G. W. Hart, Purification and characterization of an O-GlcNAc selective N-acetyl-beta-D-glucosaminidase from rat spleen cytosol. J. Biol. Chem. 269, 19321–19330 (1994).

10. Y. Gao, L. Wells, F. I. Comer, G. J. Parker, G. W. Hart, Dynamic O-glycosylation of nuclear and cytosolic proteins: cloning and characterization of a neutral, cytosolic beta-N-acetylglucosaminidase from human brain. J Biol Chem 276, 9838–9845 (2001).

11. I. Han, E. S. Oh, J. E. Kudlow, Responsiveness of the state of O-linked N-acetylglucosamine modification of nuclear pore protein p62 to the extracellular glucose concentration. Biochem. J. 350, 109–114 (2000).

12. K. Liu, A. J. Paterson, E. Chin, J. E. Kudlow, Glucose stimulates protein modification by O-linked GlcNAc in pancreatic β cells: Linkage of O-linked GlcNAc to β cell death. Proc. Natl. Acad. Sci. 97, 2820–2825 (2000).

13. N. E. Zachara, et al., Dynamic O-GlcNAc Modification of Nucleocytoplasmic Proteins in Response to Stress A SURVIVAL RESPONSE OF MAMMALIAN CELLS. J. Biol. Chem. 279, 30133–30142 (2004).

14. N. M. Akella, et al., O-GlcNAc Transferase Regulates Cancer Stem-like Potential of Breast Cancer Cells. Mol. Cancer Res. MCR 18, 585–598 (2020).

15. C. M. Ferrer, et al., O-GlcNAcylation Regulates Cancer Metabolism and Survival Stress Signaling via Regulation of the HIF-1 Pathway. Mol. Cell 54, 820–831 (2014).

16. C. M. Ferrer, V. L. Sodi, M. J. Reginato, O-GlcNAcylation in Cancer Biology: Linking Metabolism and Signaling. J. Mol. Biol. 428, 3282–3294 (2016).

17. H. M. Itkonen, et al., O-GlcNAc Transferase Integrates Metabolic Pathways to Regulate the Stability of c-MYC in Human Prostate Cancer Cells. Cancer Res. 73, 5277–5287 (2013).

18. M.-D. Li, et al., Adipocyte OGT governs diet-induced hyperphagia and obesity. Nat. Commun. 9, 1–12 (2018).

19. H. Shi, et al., Skeletal muscle O-GlcNAc transferase is important for muscle energy homeostasis and whole-body insulin sensitivity. Mol. Metab. 11, 160–177 (2018).

20. K. Vaidyanathan, L. Wells, Multiple Tissue-specific Roles for the O-GlcNAc Post-translational Modification in the Induction of and Complications Arising from Type II Diabetes. J. Biol. Chem. 289, 34466–34471 (2014).

21. Y. Yang, et al., O-GlcNAc transferase inhibits visceral fat lipolysis and promotes diet-induced obesity. Nat. Commun. 11, 181 (2020).

22. M. R. Bond, J. A. Hanover, A little sugar goes a long way: The cell biology of O-GlcNAc. J. Cell Biol. 208, 869–880 (2015).

23. G. W. Hart, C. Slawson, G. Ramirez-Correa, O. Lagerlof, Cross Talk Between O-GlcNAcylation and Phosphorylation: Roles in Signaling, Transcription, and Chronic Disease. Annu. Rev. Biochem. 80, 825–858 (2011).

24. M. B. Lazarus, et al., HCF-1 is cleaved in the active site of O-GlcNAc transferase. Science 342, 1235–1239 (2013).

25. J. Janetzko, S. A. Trauger, M. B. Lazarus, S. Walker, How the glycosyltransferase OGT catalyzes amide bond cleavage. Nat. Chem. Biol. 12, 899–901 (2016).

26. F. Capotosti, et al., O-GlcNAc Transferase Catalyzes Site-Specific Proteolysis of HCF-1. Cell 144, 376–388 (2011).

27. E. Julien, W. Herr, Proteolytic processing is necessary to separate and ensure proper cell growth and cytokinesis functions of HCF-1. EMBO J. 22, 2360–2369 (2003).

28. E. Julien, W. Herr, A switch in mitotic histone H4 lysine 20 methylation status is linked to M phase defects upon loss of HCF-1. Mol. Cell 14, 713–725 (2004).

29. S. Daou, et al., Crosstalk between O-GlcNAcylation and proteolytic cleavage regulates the host cell factor-1 maturation pathway. Proc. Natl. Acad. Sci. 108, 2747–2752 (2011).

30. W. D. Cheung, G. W. Hart, AMP-activated protein kinase and p38 MAPK activate O-GlcNAcylation of neuronal proteins during glucose deprivation. J Biol Chem 283, 13009–13020 (2008).

31. W. D. Cheung, K. Sakabe, M. P. Housley, W. B. Dias, G. W. Hart, O-linked beta-N-acetylglucosaminyltransferase substrate specificity is regulated by myosin phosphatase targeting and other interacting proteins. J Biol Chem 283, 33935–33941 (2008).

32. E. A. Lane, et al., HCF-1 Regulates De Novo Lipogenesis through a Nutrient-Sensitive Complex with ChREBP. Mol. Cell 75, 357–371.e7 (2019).

33. H.-B. Ruan, et al., O-GlcNAc transferase/host cell factor C1 complex regulates gluconeogenesis by modulating PGC-1α stability. Cell Metab. 16, 226–237 (2012).

34. H. M. Stephen, J. L. Praissman, L. Wells, Generation of an Unbiased Interactome for the Tetratricopeptide Repeat Domain of O-GlcNAc Transferase Indicates a Role for the Enzyme in Intellectual Disability. bioRxiv, 2020.07.30.229930 (2020).

35. D. Mariappa, et al., Dual functionality of O-GlcNAc transferase is required for Drosophila development. Open Biol. 5 (2015).

36. S. J. Urso, M. Comly, J. A. Hanover, T. Lamitina, The O-GlcNAc transferase OGT is a conserved and essential regulator of the cellular and organismal response to hypertonic stress. PLOS Genet. 16, e1008821 (2020).

37. A. C. Giles, et al., A complex containing the O-GlcNAc transferase OGT-1 and the ubiquitin ligase EEL-1 regulates GABA neuron function. J. Biol. Chem. 294, 6843–6856 (2019).

38. J. A. Hanover, et al., A Caenorhabditis elegans model of insulin resistance: altered macronutrient storage and dauer formation in an OGT-1 knockout. Proc. Natl. Acad. Sci. U. S. A. 102, 11266–11271 (2005).

39. H. Liu, et al., Inhibition of E-cadherin/catenin complex formation by O-linked N-acetylglucosamine transferase is partially independent of its catalytic activity. Mol. Med. Rep. 13, 1851–1860 (2016).

40. Z. Kazemi, H. Chang, S. Haserodt, C. McKen, N. E. Zachara, O-Linked β-N-acetylglucosamine (O-GlcNAc) Regulates Stress-induced Heat Shock Protein Expression in a GSK-3β-dependent Manner. J. Biol. Chem. 285, 39096–39107 (2010).

41. E. Campeau, et al., A Versatile Viral System for Expression and Depletion of Proteins in Mammalian Cells. PLoS ONE 4, e6529 (2009).

42. D. Shcherbo, et al., Far-red fluorescent tags for protein imaging in living tissues. Biochem. J. 418, 567–574 (2009).

43. A. Ramezani, R. G. Hawley, Strategies to Insulate Lentiviral Vector-Expressed Transgenes. Methods Mol. Biol. Clifton Nj 614, 77–100 (2010).

44. Z.-W. Tan, et al., O-GlcNAc regulates gene expression by controlling detained intron splicing. Nucleic Acids Res. (2020) https:/doi.org/10.1093/nar/gkaa263 (April 24, 2020).

45. G. V. Chaitanya, J. S. Alexander, P. P. Babu, PARP-1 cleavage fragments: signatures of cell-death proteases in neurodegeneration. Cell Commun. Signal. CCS 8, 31 (2010).

46. S. Gobeil, C. C. Boucher, D. Nadeau, G. G. Poirier, Characterization of the necrotic cleavage of poly(ADP-ribose) polymerase (PARP-1): implication of lysosomal proteases. Cell Death Differ. 8, 588–594 (2001).

47. Y. Liu, et al., Suppression of OGT by microRNA24 reduces FOXA1 stability and prevents breast cancer cells invasion. Biochem. Biophys. Res. Commun. 487, 755–762 (2017).

48. S.-K. Park, et al., A Conserved Splicing Silencer Dynamically Regulates O-GlcNAc Transferase Intron Retention and O-GlcNAc Homeostasis. Cell Rep. 20, 1088–1099 (2017).

49. K. Qian, et al., Transcriptional regulation of O-GlcNAc homeostasis is disrupted in pancreatic cancer. J. Biol. Chem. 293, 13989–14000 (2018).

50. V. Kapuria, et al., Proteolysis of HCF-1 by Ser/Thr glycosylation-incompetent O-GlcNAc transferase:UDP-GlcNAc complexes. Genes Dev. 30, 960–972 (2016).

51. M. B. Lazarus, Y. Nam, J. Jiang, P. Sliz, S. Walker, Structure of human O-GlcNAc transferase and its complex with a peptide substrate. Nature 469, 564–567 (2011).

52. M. B. Lazarus, et al., Structural snapshots of the reaction coordinate for O-GlcNAc transferase. Nat. Chem. Biol. 8, 966–968 (2012).

53. Z. G. Levine, et al., O-GlcNAc Transferase Recognizes Protein Substrates Using an Asparagine Ladder in the Tetratricopeptide Repeat (TPR) Superhelix. J. Am. Chem. Soc. 140, 3510–3513 (2018).

54. M. Schimpl, et al., O-GlcNAc transferase invokes nucleotide sugar pyrophosphate participation in catalysis. Nat. Chem. Biol. 8, 969–974 (2012).

55. A. Loonstra, et al., Growth inhibition and DNA damage induced by Cre recombinase in mammalian cells. Proc. Natl. Acad. Sci. U. S. A. 98, 9209–9214 (2001).

56. B. Nabet, et al., The dTAG system for immediate and target-specific protein degradation. Nat. Chem. Biol. 14, 431 (2018).

57. M. A. Erb, et al., Transcription control by the ENL YEATS domain in acute leukaemia. Nature 543, 270–274 (2017).

58. H.-T. Huang, et al., MELK is not necessary for the proliferation of basal-like breast cancer cells. eLife 6, e26693 (2017).

59. G. C. McAlister, et al., Increasing the multiplexing capacity of TMT using reporter ion isotopologues with isobaric masses. Anal. Chem. 84, 7469–7478 (2012).

60. G. C. McAlister, et al., MultiNotch MS3 enables accurate, sensitive, and multiplexed detection of differential expression across cancer cell line proteomes. Anal. Chem. 86, 7150–7158 (2014).

61. J. Navarrete-Perea, Q. Yu, S. P. Gygi, J. A. Paulo, Streamlined Tandem Mass Tag (SL-TMT) Protocol: An Efficient Strategy for Quantitative (Phospho)proteome Profiling Using Tandem Mass Tag-Synchronous Precursor Selection-MS3. J. Proteome Res. 17, 2226–2236 (2018).

62. E. P. Roquemore, et al., Vertebrate lens alpha-crystallins are modified by O-linked N-acetylglucosamine. J. Biol. Chem. 267, 555–563 (1992).

63. C. M. Woo, et al., Mapping and Quantification of Over 2000 O-linked Glycopeptides in Activated Human T Cells with Isotope-Targeted Glycoproteomics (Isotag). Mol. Cell. Proteomics MCP 17, 764–775 (2018).

64. B. Jassal, et al., The reactome pathway knowledgebase. Nucleic Acids Res. 48, D498–D503 (2020).

65. M. Kanehisa, S. Goto, KEGG: kyoto encyclopedia of genes and genomes. Nucleic Acids Res. 28, 27–30 (2000).

66. A. Subramanian, et al., Gene set enrichment analysis: A knowledge-based approach for interpreting genome-wide expression profiles. Proc. Natl. Acad. Sci. 102, 15545–15550 (2005).

67. S. E. S. Martin, et al., Structure-Based Evolution of Low Nanomolar O-GlcNAc Transferase Inhibitors. J. Am. Chem. Soc. 140, 13542–13545 (2018).

68. R. P. Taylor, et al., Glucose deprivation stimulates O-GlcNAc modification of proteins through up-regulation of O-linked N-acetylglucosaminyltransferase. J. Biol. Chem. 283, 6050–6057 (2008).

69. X. Li, et al., O-GlcNAcylation of core components of the translation initiation machinery regulates protein synthesis. Proc. Natl. Acad. Sci. U. S. A. 116, 7857–7866 (2019).

70. T. Ohn, N. Kedersha, T. Hickman, S. Tisdale, P. Anderson, A functional RNAi screen links O-GlcNAc modification of ribosomal proteins to stress granule and processing body assembly. Nat. Cell Biol. 10, 1224–1231 (2008).

71. Q. Zeidan, Z. Wang, A. De Maio, G. W. Hart, O-GlcNAc cycling enzymes associate with the translational machinery and modify core ribosomal proteins. Mol. Biol. Cell 21, 1922–1936 (2010).

72. K. E. Pyo, et al., ULK1 O-GlcNAcylation Is Crucial for Activating VPS34 via ATG14L during Autophagy Initiation. Cell Rep. 25, 2878–2890.e4 (2018).

73. H.-B. Ruan, et al., Calcium-dependent O-GlcNAc signaling drives liver autophagy in adaptation to starvation. Genes Dev. 31, 1655–1665 (2017).

74. P. Wang, J. A. Hanover, Nutrient-driven O-GlcNAc cycling influences autophagic flux and neurodegenerative proteotoxicity. Autophagy 9, 604–606 (2013).

75. E. P. Tan, et al., Altering O-Linked β-N-Acetylglucosamine Cycling Disrupts Mitochondrial Function. J. Biol. Chem. 289, 14719–14730 (2014).

76. W. Liu, et al., PHF8 mediates histone H4 lysine 20 demethylation events involved in cell cycle progression. Nature 466, 508–512 (2010).

77. J. Wysocka, M. P. Myers, C. D. Laherty, R. N. Eisenman, W. Herr, Human Sin3 deacetylase and trithorax-related Set1/Ash2 histone H3-K4 methyltransferase are tethered together selectively by the cell-proliferation factor HCF-1. Genes Dev. 17, 896–911 (2003).

78. S. Minocha, et al., Rapid Recapitulation of Nonalcoholic Steatohepatitis upon Loss of Host Cell Factor 1 Function in Mouse Hepatocytes. Mol. Cell. Biol. 39 (2019).

79. T. M. Kristie, Y. Liang, J. L. Vogel, Control of α-herpesvirus IE gene expression by HCF-1 coupled chromatin modification activities. Biochim. Biophys. Acta BBA - Gene Regul. Mech. 1799, 257–265 (2010).

80. M. L. Hancock, et al., Insulin Receptor Associates with Promoters Genome-wide and Regulates Gene Expression. Cell 177, 722–736.e22 (2019).

81. J. L. Vogel, T. M. Kristie, Site-specific proteolysis of the transcriptional coactivator HCF-1 can regulate its interaction with protein cofactors. Proc. Natl. Acad. Sci. U. S. A. 103, 6817–6822 (2006).

82. L. Franco-Serrano, et al., Multifunctional Proteins: Involvement in Human Diseases and Targets of Current Drugs. Protein J. 37, 444–453 (2018).

83. G. M. Burslem, C. M. Crews, Proteolysis-Targeting Chimeras as Therapeutics and Tools for Biological Discovery. Cell 181, 102–114 (2020).

84. H. M. Itkonen, et al., Inhibition of O-GlcNAc transferase renders prostate cancer cells dependent on CDK9. Mol. Cancer Res. MCR (2020) https:/doi.org/10.1158/1541-7786.MCR-20-0339.

