## Supplemental Text, Figures, and Methods for "Mammalian Cell Proliferation Requires Noncatalytic Functions of O-GlcNAc Transferase"

### *Supplementary Materials and Methods*

#### *Cell Lines*

HEK 293T (ATCC CRL-3216) cells were grown in DMEM (low glucose with pyruvate, Glutamax; ThermoFisher 10567-022) + 10% fetal bovine serum (FBS; ThermoFisher 10437-028) in a 5% $CO_2$ humidified incubator.

Inducible *Ogt* knockout MEFs (1) were grown in DMEM + 10% FBS in a 5% $CO_2$ humidified incubator. To ensure high efficiency of knockout, single clones of this MEF line were isolated by limiting dilution cloning. Incucyte growth curve experiments were performed in the presence of 1x Penicillin/Streptomycin (Corning 30-002-CI). All cell experiments used cell lines at fewer than 10 passages from receipt or from clonal isolation in newly-derived cell lines. Single clones of FKBP12$^{F36V}$-OGT MEFs were isolated by limiting dilution. Cell lines were regularly checked for mycoplasma using MycoAlert Detection Kit (Lonza LT07-118).

HeLa S3 (ATCC CCL-2.2) for cell extract preparation were grown in DMEM + 10% FBS in a 5% $CO_2$ incubator as previously described (2).

#### Lentivirus Production

HEK293T cells were plated on poly-D-lysine (Sigma Aldrich P6407) coated 10 cm plates at $7\times10^6$ cells per plate in and allowed to attach to plate overnight. Per 10 cm plate, 1.5 $\mu$g pMD2.G, 3.5 $\mu$g psPAX2, and 9.5 $\mu$g OGT lentiviral plasmid were mixed in serum free DMEM (100 $\mu$L final volume). 60 $\mu$g polyethyleneimine solution (PEI, Polyscience inc. cat 23966; made up at 1.0 mg/mL in distilled water, pH-adjusted with HCl to pH 7.0 and filter sterilized) was added and mixed thoroughly. Media on cells in 10 cm dish was changed to 10 mL fresh DMEM + 10% FBS. After 10–15 minutes, DNA/PEI mixture was added dropwise with gentle mixing to cells, and cells were returned to incubator.

After overnight incubation, media was removed and replaced with 5 mL DMEM. At 48 hours after transfection, media was collected, spun at 500 x g in centrifuge, and supernatant was either aliquoted in 2 mL aliquots and flash frozen or used directly to infect cells. A second 5 mL media portion was added, and a second batch of virus collected at 72 hours after transfection.

#### Lentiviral Infection of MEF Cells

MEF cells were plated at $2 \times 10^5$ cells/well in 6 well plates. Cells were allowed to attach to plate overnight. Swinging-bucket centrifuge was warmed to 30 °C. 1.6 $\mu$L of 10 mg/mL polybrene was added to 2 mL viral aliquots and mixed. Media was removed from cells and 2 mL viral aliquot was added to each well. Plate was placed in centrifuge and spun at 1000 x g for 90 minutes. 1 mL media was removed and DMEM + 10% FBS with 8 $\mu$g/mL polybrene was added, and cells were returned to 5% $CO_2$ incubator overnight. Media was replaced with fresh DMEM + 10% FBS after overnight incubation. At 48 hours after infection, selection antibiotic (1.0 mg/mL Neomycin for non-mKate2 viruses, including $FKBP12^{F36V}$-OGT; 2.0 $\mu$g/mL Puromycin for mKate2-OGT constructs) was added. Cells were maintained and expanded in antibiotic for at least one week before removing selective media. Selected cells were either frozen, sorted for fluorescence, or passaged at least once before subjecting to assays.

#### Fluorescent Sorting of mKate2-OGT Cells

MEF cells were trypsinized with TrypLE and re-suspended as single-cell suspensions, which were passed through a 40 $\mu$m strainer. The base cell line without mKate-OGT lentivirus was collected as a negative control. Cells were sorted on either a BD FACSAria (594 nm laser, 610/20 bandpass filter) or MoFlo Astrios EQ (592 nm laser, 620/29 bandpass filter) at the Harvard Medical School Department of Immunology Flow Cytometry Core. Gates were set at a level that would capture fewer than 0.1% of cells in the negative control, providing clear expression of mKate2-OGT over background.

#### MEF Cell Line Isolation

To ensure high-efficiency OGT knockout in drug treatments, single clones were isolated by limiting dilution of both $FKBP12^{F36V}$-OGT and initial Cre-based inducible *Ogt* null cells. Cre-based inducible *Ogt* null cell clones were tested by live cell imaging-based growth curve analysis for efficient knockout before use in further assays. $FKBP12^{F36V}$-OGT clones were isolated by limiting dilution two weeks after 4HT treatment, and tested by western blot for loss of endogenous OGT and robust $FKBP12^{F36V}$-OGT expression (Figure 3B).

#### Plasmid Cloning and Molecular Biology

All entry vector plasmids were amplified in Stellar *E. coli* cloning strain in LB grown at 37 °C; all lentiviral plasmids were maintained in NEBStable cloning strain (New England Biolabs #C3040) grown at 30 °C. Transformation was carried out according to manufacturers protocols. All individual plasmids were confirmed by Sanger sequencing of complete insert and plasmid junction before use. All oligonucleotides and gBlocks were purchased from Integrated DNA Technologies.

Human OGT coding sequence was PCR amplified with KOD polymerase (EMD Millipore 71086-4) from cDNA (1036 amino acid isoform, ENTREZ ID: NM_181673.3) with primers containing BamHI and XhoI restriction sites at the 5' and 3' ends respectively. Vector pENTR1A (no ccdB) and PCR product were restriction digested and vector was treated with alkaline phosphatase as per manufacturers protocol. Both PCR product and vector were gel purified. T4 ligase was used to ligate vector and insert as per manufacturer's protocol.

All point mutants of human OGT were introduced using Q5 site directed mutagenesis (New England Biolabs #E0554) as per manufacturers protocol. 5N5A mutant was prepared by PCR-linearization using Q5 DNA polymerase (New England Biolabs #M0491). 5N5A mutations were inserted using isothermal assembly(3) from a gBlock containing the desired mutations using the Gibson Assembly Master Mix as per the manufacturers instructions (New England Biolabs #E2611). The same overall process was used to insert mKate2 into pENTR1A OGT vectors.

To construct FKBP12$^{F36V}$-OGT vectors, Q5 site-directed mutagenesis was used to generate N-terminal "open" human OGT pENTR1A vectors, which were inserted into pLEX_305-N-dTAG using LR Clonase II according to manufacturer's directions. Due to poor compatibility of this vector with the MEF cell line, the FKBP12$^{F36V}$-OGT insert was amplified with primers containing BamHI and NotI sites at the 3' and 5' ends of FKBP12$^{F36V}$-OGT respectively. pENTR1A (no ccdB) and FKBP12$^{F36V}$-OGT PCR product were digested and ligated together as described above.

For FKBP12$^{F36V}$-OGT and untagged OGT constructs, LR recombination with LR Clonase II (ThermoFisher #11791020) was used to transfer insert into pLenti PGK Neo DEST. For all mKate2-OGT variants, LR Clonase II was used to insert into pLenti PGK Puro DEST.

#### Recombinant OGT Purification

All proteins were grown in LOBSTR *E. coli* expression strain (Kerafast #EC1001), which were transformed with plasmid according to manufacturers protocol. Expression strain was grown in LB with shaking (250 rpm) to 37 °C to OD 0.8–1.0. Cultures were cooled to 16 °C for 30 minutes and expression was induced by addition of IPTG to a final concentration of 0.2 mM; expression proceeded at 16 °C for 16 hours with continued shaking at 250 rpm. Cells were then pelleted at 4000 x g, and resuspended in TBS (pH 7.4) with 0.1 mg/mL DNAse I, 0.1 mg/mL Lysozyme, and PMSF 1 mM. Resuspended cells were lysed using an Avestin Emulsiflex C3 cell disruptor 3 times at 15000 psi. Lysate was clarified by centrifiguation at 14000 x g for 20 minutes at 4 °C.

Imidazole pH 8.0 was added to clarified lysate to final concentration of 40 mM, and 1 mL packed Ni-NTA Agarose (pre-equilibrated with TBS pH 7.4 40 mM imidazole) was added per liter culture. Resin was incubated with lysate with mixing via rotation for 1 hour, then flow through collected. Resin was washed with 10 column volumes TBS pH 7.4 50 mM imidazole, then protein was eluted using 4 column volumes TBS pH 7.4 250 mM imidazole. Elution fraction were supplemented with tris(hydroxypropyl)phosphine (THP) to 1 mM. Elution fractions were concentrated using 100 kDa Amicon spin concentrators. Protein was further purified via size-exlusion chromatography using a Superdex 200 increase 10/300 GL column, concentrated via 100 kDa Amicon spin concentrator, aliquoted, and flash frozen. All batches were checked for purity by Coomassie staining of SDS-PAGE. Upon thawing, protein aliquots were spun at 21,000 x g for 20 minutes at 4 °C, and supernatant collected to remove any aggregate. Concentration was assessed by $A_{280}$ on a Nanodrop using an extinction coefficient of 117580 $M^{-1}cm^{-1}$.

### HeLa Cell Extract Preparation

Cell extracts were prepared as previously described (Levine et al.). Briefly, a 1 L spinner culture of HeLa S3 cells were grown to $2x10^6$ cells/mL, pelleted, and flash frozen. Upon thawing on ice, cells were resuspended in swelling buffer (pH 7.5, 40 mM HEPES, 5 mM KCl, 1.5 mM $MgCl_2$, 1 mM DTT, 1 EDTA-free protease inhibitor cocktail tablet (Roche) per 25 mL) of approximately 80% the volume of the pelleted cell mass. Energy mix (7.5 mM creatine phosphate, 1 mM ATP, 0.1 mM EGTA, pH 7.7, 1 mM $MgCl_2$) was added, and the resuspended cell mass was stirred at 4 °C under 1000 psi pressure nitrogen gas for 30 minutes, after which the pressure was release and the cellular lysate collected, followed by a second round of pressurization and pressure release to ensure complete lysis. Precipitates were pelleted at 15000 rcf for 30 minutes. Cell extracts were aliquoted and flash frozen.

### HeLa Cell Extract Glycosylation

HeLa cell extracts were thawed on ice and centrifuged at 4 ˚C for 20 minutes at 45,000 rpm (Beckman Coulter TLA120.2 rotor) to pellet any remaining precipitated debris. Following centrifugation, the HeLa cell extracts were aliquoted to a final volume of 40 µL. The aliquots were incubated with 1 mM of UDP-GlcNAc (Promega V7071) and equal volumes of reaction buffer (20 mM Tris, pH 7.4, 150 mM NaCl, 20 mM $MgCl_2$, 1 mM trishydroxypropylphosphine) or 2.5 mM OGT at 37 ˚C. At 1 minute, 5 minutes, and 15 minutes, 10 µL aliquots were taken from the reaction and quenched with 10 µL 2X Laemmli loading buffer. The samples were boiled at 95 ˚C for 10 minutes and run on a 4-20% TGX SDS-PAGE gel (BioRad) at 180 V for 1.5 hours. The gels were transferred to nitrocellulose membranes and analyzed by western blot with the anti-O- GlcNAc (CTD110.6, Cell Signaling), anti-OGT (D1D8Q, Cell Signaling), and anti-GAPDH (9484, Abcam) antibodies.

### OGT knockout in Cre/Lox System

$Ogt^{f/Y}$ MEF cells were plated 24 hours before treatment in DMEM + 10% FBS. Media was changed to DMEM + 10% FBS + 500 nM 4HT (diluted from 10 mM stock in ethanol) at time zero. After 24 hours, media was replaced with DMEM + 10% FBS. Media was changed daily thereafter until cells were collected for lysis.

### OGT Degradation with dTAG-13

$FKBP12^{F36V}$-OGT MEF cells were plated 24 hours before treatment in DMEM + 10% FBS. Media was changed to DMEM + 10% FBS + 500 nM dTAG-13 (diluted from 10 mM stock in DMSO) at time zero. Every 24 hours media was removed and replaced with DMEM + 10% FBS with 500 nM dTAG-13 until cell collection.

### MEF Growth Curves

All cells were grown in DMEM supplemented with 10% FBS and 1x Pen/Strep antibiotics. At the day of plating, cells were diluted to 5000 cells/mL and plated at 100 $\mu$L per well in a 96 well plate . The outer edge of the 96 well plate was filled with 200 $\mu$L PBS to maintain hydration of all wells containing cells.

*Cre-based OGT Deletion*

At 24 hours after plating, media was replaced with 100 $\mu$L of DMEM containing 500 nM (Z)-4-hydroxytamoxifen (4HT; diluted from 10 mM stock in ethanol). Within 1 hour of initial treatment, cells were placed in Essen Incucyte Zoom for imaging every 4 hours. At 24 hours and once a day thereafter, cells were removed and media changed to DMEM with FBS and Pen/Strep; no 4HT was included after the first 24 hours. If a cell line had reached 85% confluence or higher at time of media change, cells were split 1:3 and replated for the length of the experiment. For each data point, two independent clones were grown with 3 technical replicates each, for a total of 6 data points.

*FKBP12$^{F36V}$-OGT degradation*

At 24 hours after plating, media was replaced with 100 $\mu$L of DMEM containing 500 nM dTAG-13 (diluted from 10 mM stock in DMSO). Within 1 hours of initial treatment, cells were placed in Essen Incucyte Zoom for imaging every 4 hours. At 24 hours and once a day thereafter, cells were removed and media changed to fresh DMEM with 500 nM dTAG-13. If a cell line had reached 85% confluence or higher at time of media change, cells were split 1:3 and replated for the length of the experiment. For each data point, two independent clones were grown with 3 technical replicates each, for a total of 6 data points.

*Analysis of growth curves*

Confluence values were obtained using the Basic Analyzer tool of the Incucyte Zoom Software version 2016b.

For each well, all confluence values were divided by the initial confluence value at the first time point. The $\log_2$ transform of this confluence ratio is obtained to determine number of doublings per well. Confluence values after each split are raised to be equal to the previous measurement, thus measuring number of doublings after the split and adding to those prior. Average and standard error of the mean were calculated using these transformed values. All analysis was carried out using custom python scripts.

### Immunoblot Analysis of OGT Activity In Cells

Cells treated as described above were grown in 10 cm tissue culture-treated plates, then washed 2x with PBS, scraped into PBS, pelleted, and flash frozen. Cells were plated day before treatment with 4HT or dTAG-13 at levels to allow 90% confluence upon harvest based upon growth rates.

To frozen cell pellets, RIPA buffer was added with 50 $\mu$M PUGNAc (Sigma Aldrich, A7229), 1x PhosSTOP (Sigma-Aldrich #4906845001), and 1x EDTA-Free HALT protease (Sigma-Aldrich 11873580001). Cells were triturated in RIPA on ice until thawed and full dissolved, then spun at 4 °C at 20,000 x g for 20 minutes. The supernatant was removed, and 10% SDS was added to bring final concentration to 2%.

BCA Assay was used to determine protein concentration, and samples were mixed with Laemmli Blue loading dye (6x), boiled for 5 minutes at 95 °C and run on 4–20% tris-glycine gels (Criterion TGX, Bio-Rad). Gels were transferred to Immobilon-FL PVDF using the TransBlot Turbo System (Bio-Rad) on the preset High MW program.

Membranes were blocked with SuperBlock (TBS; ThermoFisher #37535) for 1 hour rocking at room temperature. Membranes were then incubated with primary antibody (OGA/MGEA5- Sigma Aldrich #HPA036141; O-GlcNAc (RL2) Abcam #ab2739; HCF-1 N terminus Cell Signaling Technology # 69690S; PARP1 Cell Signaling Technology #9542S; all 1:1000 dilution) diluted in SuperBlock T20 (ThermoFisher #37536) overnight at at 4 °C with rocking; OGT (Cell Signaling Technology and actin were co-incubated (at 1:1000 dilution each).

After overnight incubation, membranes were washed 3x10 minutes with TBS-T at room temperature and incubated for 1 hour with secondary antibody. OGT (Cell Signaling Technology #24083) and actin (Cell Signaling Teechnology #3700S) were analyzed by fluorescent secondaries Goat Anti-Rabbit IgG IRDye 800CW and Donkey Anti-Mouse IgG IRDye 680RD (1:10,000 dilution each; LI-COR Biosciences #926-32211, #926-68072) in Superblock T20 with 0.02% SDS. All others were incubated with appropriate HRP-conjugated secondary antibody in Superblock T20 (1:2000 dilution).

Blots were washed 3 x 10 minutes TBS-T. HRP-blots were imaged using Thermo Scientific™ SuperSignal™ West Pico PLUS Chemiluminescent Substrate according to manufacturer's instructions using an Azure C600 Imager. OGT/actin blots were washed further in TBS pH 7.4 before imaging on near-IR settings with Azure C600 Imager.

### Quantitative Proteomics Analysis of Protein Expression

*Sample Preparation and Liquid Chromatography/Mass Spectrometry*

FKBP12$^{F36V}$-OGT MEF cell lines were plated in 10 cm dishes and allowed to settle onto plate overnight. The next day, cells were either collected or treated with dTAG-13 500 nM in fresh DMEM + 10% FBS. At 24 hours after treatment, day one samples were collected, while fresh media with 500 nM dTAG-13 were introduced to 48 hour samples, which were collected at 48 hours after initial treatment. Collection was performed by removing cells from dish by TrypLE trypsinization, quenching with cold DMEM + 10 % FBS, pelleting at 500 x g, washing with ice cold PBS, and pelleting again before flash-freezing in liquid nitrogen and storing at -80 until all samples were ready. For each cell type, two biological replicates, consisting of mKate2-OGT lentiviral infections of two different FKBP12$^{F36V}$-OGT clones, were collected.

Upon collection of all samples, samples were lysed by resuspension of frozen sample in 100–200 $\mu$L of proteomics lysis buffer (50 mM Tris pH 8.8, 150 mM NaCl, 2% SDS, phosStop phosphatase inhibitors, HALT EDTA-free protease inhibitor, 5 mM DTT). Samples were passed through a 26-gauge needle 10x to shear DNA then spun at 21000 x g for 20 minutes at 12 °C.

The supernatant was incubated at 60 °C for 45 minutes, then allowed to cool to room temperature. Iodoacetamide (0.5 M stock solution made fresh in distilled water) was added to a final concentration of 14 mM and sample was incubated in the dark for 45 minutes. Protein concentration of samples was measured using BCA assay.

100 $\mu$g of sample was aliquoted out. If more concentrated, samples were diluted to 1 $\mu$g/$\mu$L with TBS + 2% SDS. 3 volumes methanol were added, followed by 100 $\mu$L chloroform and 250 $\mu$L water, at which point samples were vortexed and centrifuged at 21000 x g for 1 minute at room temperature. Leaving the precipitated protein undisturbed, removed solvent and added 1 mL methanol to wash, then centrifuged 21000 x g 1 minute at room temperature.

Solvent was removed, samples were processed using the SL-TMT protocol (4). In brief, protein pellets were resuspended in 200 mM EPPS pH 8.5 to 1 mg/ml. Proteins were disgested with LysC (substrate:enzyme = 100) overnight at 37$^{o}$C and then with Trypsin (substrate:enzyme = 100) for 6 hours at 37$^{o}$C. 3.3 $\mu$L was removed from each sample and mixed in a 31$^{st}$ tube to generate our mixed normalization sample. The resulting 31 peptide solutions were then labelled with TMT 10/11-plex reagents (Thermo Scientific) for 1.5 hour at room temperature. Reactions were stopped by addition of 5% hydroxylamine for 30 minutes. Equal amount of peptide samples were combined, dried by vacuum centrifugation and desalted on a Waters C18 solid phase extraction Sep-Pak. TMT-labeled peptide samples were fractionated via BPRP HPLC to 96 fractions, which were consolidated to 12. These fractions were subsequently acidified with 1% formic acid, vacuum centrifuged to near dryness and deslated with C18 stagetips. Dried peptides were resuspended in 5% acetronitrile/ 5% formic acid for LC-MS/MS processing.

Our mass spectrometry data were collected using an Orbitrap Fusion mass spectrometer (ThermoFisher Scientific, San Jose, CA) coupled to a Proxeon EASY-nLC 1000 liquid chromatography (LC) pump (Thermo Fisher Scientific). Peptides were separated on a 100 µm inner diameter microcapillary column packed with 35 cm of Accucore C18 resin (2.6 µm, 150 Å, ThermoFisher). For each analysis, we loaded ~2 µg onto the column. Separation was in-line with the mass spectrometer and was performed using a 3 hr gradient of 6 to 26% acetonitrile in 0.125% formic acid at a flow rate of ~450 nL/min. Each analysis used a TMT-based TMT method, which has been shown to reduce ion interference compared to MS2 quantification (5). The scan sequence began with an MS1 spectrum (Orbitrap analysis; resolution 120,000; mass range 400−1400 *m/z*; automatic gain control (AGC) target $5 \times 10^5$; maximum injection time 100 ms). Precursors for MS2/MS3 analysis were selected using a Top10 method. MS2 analysis consisted of collision-induced dissociation (CID); AGC $2.0\times 10^4$; normalized collision energy (NCE) 35; maximum injection time 120 ms; and isolation window of 0.4 Da. Following acquisition of each MS2 spectrum, we collected an MS3 spectrum using a recently described method in which multiple MS2 fragment ions were captured in the MS3 precursor population using isolation waveforms with multiple frequency notches (6). MS3 precursors were fragmented by high energy collision-induced dissociation (HCD) and analyzed using the Orbitrap (NCE 65; AGC $1.5 \times 10^5$; maximum injection time 150 ms, resolution was 50,000 at 400 Th, isolation window 0.7 Da).

### Quantification and Statistical Analysis

#### Analysis of OGT Conservation in gnomAD

Genome-wide data on human gene conservation was downloaded from the gnomAD database of >140,000 human exome and full genomes (7). Ratio of observed missense mutations to expected missense mutations were plotted as a histogram for all human genes; ratios were used as calculated based on background mutation rate (7). Histograms were truncated at 2.0 to not include hyper-variable genes. Vertical lines were added at correct missense mutation ratio for genes of interest. All analysis was performed using custom python scripts. Marked glycosyltransferase genes were those annotated in (8).

### Quantitative Proteomic Data Analysis

Mass spectra were processed using a SEQUEST-based pipeline (9–11). Spectra were converted to mzXML using a modified version of ReAdW.exe. Database searching included all entries from the human UniProt database. This database was concatenated with one composed of all protein sequences in the reversed order. Searches were performed using a 50 ppm precursor ion tolerance for total protein level analysis. The product ion tolerance was set to 0.9 Da. These wide mass tolerance windows were chosen to maximize sensitivity in conjunction with Sequest searches and linear discriminant analysis (12, 13). TMT tags on lysine residues and peptide N termini (+229.163 Da) and carbamidomethylation of cysteine residues (+57.021 Da) were set as static modifications, while oxidation of methionine residues (+15.995 Da) was set as a variable modification.

Peptide-spectrum matches (PSMs) were adjusted to a 1% false discovery rate (FDR) (14, 15). PSM filtering was performed using a linear discriminant analysis, while considering the following parameters: XCorr, ΔCn, missed cleavages, peptide length, charge state, and precursor mass accuracy. For TMT-based reporter ion quantitation, we extracted the signal-to-noise (S:N) ratio for each TMT channel and found the closest matching centroid to the expected mass of the TMT reporter ion. PSMs were identified, quantified, and collapsed to a 1% peptide false discovery rate (FDR) and then collapsed further to a final protein-level FDR of 1%. Moreover, protein assembly was guided by principles of parsimony to produce the smallest set necessary to account for all observed peptides.

Peptide intensities were quantified by summing reporter ion counts across all matching PSMs, as described previously (6, 16). Briefly, a 0.003 Th window around the theoretical m/z of each reporter ion was scanned for ions, and the maximum intensity nearest the theoretical m/z was used. PSMs with poor quality, MS3 spectra with TMT reporter summed signal-to-noise ratio less than 100, or no MS3 spectra were excluded from quantitation, and isolation specificity ≥ 0.7 was required (16).

#### Multiple Linear Regression-Based Analysis of Protein Expression

Protein abundance for each protein on each day was divided by protein abundance in normalization channel to generate data comparable across all 3 days. For multi-day data, all proteins that were missing on any single day were removed.

For comparison between Day 0 and Days 1 or 2, we did a Welch's t-test of $\log_2$ transformed protein abundance. We suggest interpreting individual hits with caution as duplicate data leaves us unable to accurately estimate variance of individual samples for a t-test (leading to bootstrap based p-values below). However, we reasoned this method could be used to determine relative amount of protein changes between each sample. A p-value cutoff of 0.01 was used as a moderately stringent cutoff for these determinations.

All analysis was performed with custom python scripts.

*Single Day Regression*

The vector $Y$ consists of $\log_2$ protein abundances, in the order *ΔOGT* replicate 1, *ΔOGT* replicate 2, $OGT^{WT}$ replicate 1, $OGT^{WT}$ replicate 2, $OGT^{\Delta Cat}$ replicate 1, $OGT^{\Delta Cat}$ replicate 2, $OGT^{\Delta GlcNAc}$ replicate 1, $OGT^{\Delta GlcNAc}$ replicate 2, $OGT^{\Delta Clvg}$ replicate 1, $OGT^{\Delta Clvg}$ replicate 2. Ordinary linear least squares was used to fit the equation

$$Y = X\beta + \varepsilon$$

where $X$ is the transpose of the matrix shown in the inset of Figure 4B, doubled to allow replicates with a column added for the average $OGT^{WT}$ expression level $C$:

$$X = \begin{bmatrix} 1 & 1 & 1 & 1 \\ 1 & 1 & 1 & 1 \\ 0 & 0 & 0 & 1 \\ 0 & 0 & 0 & 1 \\ 1 & 1 & 0 & 1 \\ 1 & 1 & 0 & 1 \\ 1 & 0 & 0 & 1 \\ 1 & 0 & 0 & 1 \\ 0 & 1 & 0 & 1 \\ 0 & 1 & 0 & 1 \end{bmatrix}$$

and

$$\beta = \begin{bmatrix} \beta_{GlcNAc} \\ \beta_{Clvg} \\ \beta_{Ncat} \\ C \end{bmatrix}$$

This equation is solved for $\beta$ using ordinary least squares for each protein in the proteome.

In order to obtain p-values, a null distribution is generated by resampling with replacement. The same regression was run on 10 randomly re-sampled values from the $\log_2$ protein abundance obtained from the 10 cell lines for each protein and regression was performed as above to obtain null $\beta$ values. This was repeated $10^6$ times. The actual p-value was obtained as the proportion of null $\beta$ values of each type more extreme than the actual measured value; as one-sided measures were used, proportions were doubled to account for two-sided hypothesis testing. The results are shown in Dataset S3.

*Multi-Day Regression*

Multi-day regression was carried out as described above for day 2 data with the following modifications. All three variables were expanded to include the full set of $\log_2$ proteomics data normalized between days. $Y$ was a vector of length 30, with the samples ordered as described above for day 2, but with day 0 samples before day 1 samples, followed by day 2 samples. All proteins lacking data for all 3 days were removed.

$X$ and $\beta$ were as shown below (with $X$ shown as the transpose):

$$X^T = \begin{bmatrix} 1 & 1 & 0 & 0 & 1 & 1 & 1 & 1 & 0 & 0 & 0 & 0 & 0 & 0 & 0 & 0 & 0 & 0 & 0 & 0 & 0 & 0 & 0 & 0 & 0 & 0 & 0 & 0 & 0 & 0 \\ 1 & 1 & 0 & 0 & 1 & 1 & 0 & 0 & 1 & 1 & 0 & 0 & 0 & 0 & 0 & 0 & 0 & 0 & 0 & 0 & 0 & 0 & 0 & 0 & 0 & 0 & 0 & 0 & 0 & 0 \\ 1 & 1 & 0 & 0 & 0 & 0 & 0 & 0 & 0 & 0 & 0 & 0 & 0 & 0 & 0 & 0 & 0 & 0 & 0 & 0 & 0 & 0 & 0 & 0 & 0 & 0 & 0 & 0 & 0 & 0 \\ 0 & 0 & 0 & 0 & 0 & 0 & 0 & 0 & 0 & 0 & 1 & 1 & 0 & 0 & 1 & 1 & 1 & 1 & 0 & 0 & 0 & 0 & 0 & 0 & 0 & 0 & 0 & 0 & 0 & 0 \\ 0 & 0 & 0 & 0 & 0 & 0 & 0 & 0 & 0 & 0 & 1 & 1 & 0 & 0 & 1 & 1 & 0 & 0 & 1 & 1 & 0 & 0 & 0 & 0 & 0 & 0 & 0 & 0 & 0 & 0 \\ 0 & 0 & 0 & 0 & 0 & 0 & 0 & 0 & 0 & 0 & 1 & 1 & 0 & 0 & 0 & 0 & 0 & 0 & 0 & 0 & 0 & 0 & 0 & 0 & 0 & 0 & 0 & 0 & 0 & 0 \\ 0 & 0 & 0 & 0 & 0 & 0 & 0 & 0 & 0 & 0 & 0 & 0 & 0 & 0 & 0 & 0 & 0 & 0 & 0 & 0 & 1 & 1 & 0 & 0 & 1 & 1 & 1 & 1 & 0 & 0 \\ 0 & 0 & 0 & 0 & 0 & 0 & 0 & 0 & 0 & 0 & 0 & 0 & 0 & 0 & 0 & 0 & 0 & 0 & 0 & 0 & 1 & 1 & 0 & 0 & 1 & 1 & 0 & 0 & 1 & 1 \\ 0 & 0 & 0 & 0 & 0 & 0 & 0 & 0 & 0 & 0 & 0 & 0 & 0 & 0 & 0 & 0 & 0 & 0 & 0 & 0 & 1 & 1 & 0 & 0 & 0 & 0 & 0 & 0 & 0 & 0 \\ 1 & 1 & 1 & 1 & 1 & 1 & 1 & 1 & 1 & 1 & 1 & 1 & 1 & 1 & 1 & 1 & 1 & 1 & 1 & 1 & 1 & 1 & 1 & 1 & 1 & 1 & 1 & 1 & 1 & 1 \end{bmatrix}$$

$$\beta = \begin{bmatrix} \beta_{GlcNAc\_d0} \\ \beta_{Clvg\_d0} \\ \beta_{Ncat\_d0} \\ \beta_{GlcNAc\_d1} \\ \beta_{Clvg\_d1} \\ \beta_{Ncat\_d1} \\ \beta_{GlcNAc\_d2} \\ \beta_{Clvg\_d2} \\ \beta_{Ncat\_d2} \\ C \end{bmatrix}$$

Note that multi-day regression measured distance from average of $OGT^{WT}$ samples to have effect sizes represent change upon loss of a given OGT activity. Constant $C$ represents average $OGT^{WT}$ expression level. As described above, this procedure was repeated for all proteins for which data was available for all three days. P-values were obtained from resampling of all 30 values. Effect sizes and p-values for all $\beta$ coefficients are reported in Dataset S4.

### Analysis of Activity-Based Effect Sizes

For all correlation analysis between regression-derived effect sizes and pairs of cell lines, the difference in average $\log_2$ protein abundances for each protein for each pair of cells indicated in Figure S4A (representing the $\log_2$ fold change) was correlated with proteome-wide effect sizes from multi-day regression using Pearson's method. For correlations between effect sizes and changes in protein abundance in individual cell lines over time, difference in average $\log_2$ protein abundance between dTAG-13 treated cells and untreated controls was correlated with effect sizes from multi-day regression using Pearson's method.

Hits for effect sizes had an effect size ($\beta$) with absolute value greater than $\log_2(1.5)$, representing at least a 1.5-fold change, and a p-value $< 0.05$. For the heatmap in Figure 5B, proteins were clustered by hierarchical clustering using Ward's method (17) based upon pattern of effect sizes across all 3 activities for all 3 days.

All plotting was performed using custom python scripts.

#### Comparison of Effect Sizes to Known O-GlcNAc Proteins and OGT Interactors

Human genes that are orthologs of all detected proteins were determined for alignment to proteomics studies performed in humans using BioMart(18). Data on O-GlcNAc glycopeptides detected in human T-cells (Woo et al. Table S1(19)) was reduced to the set of human genes for which at least one glycopeptide was discovered. This data, along with data about OGT interactors from affinity purification-mass spectrometry (Ruan et al. Table S1 (20), manually annotated with HGNC symbols) was aligned to proteomics data on the basis of having the same gene name as a protein's human ortholog. These proteins were compared for enrichment in proteins that had a p-value of less than 0.01 on Day 2 for effect sizes for each OGT activity based upon the multiday regression analysis described above. To measure rates, the number of proteins both in the O-GlcNAc proteins or OGT interactor dataset and meeting the p-value cutoff was divided by the full set meeting the p-value cutoff; this was compared to the total number of O-GlcNAc proteins (1250) or OGT interactors (314) divided by 7220, the total number of proteins analyzed in our study. To test whether enrichment was statistically significant, we used a permutation test with n=10000 permutations. For each permutation, we randomly selected (without replacement) the same number of proteins as in the actual dataset (1250 or 314) and tested how many met the p-value cutoffs for each OGT activity at day 2 divided by the total number of proteins meeting that p-value cutoff. After collecting 10000 such ratios for each OGT activity and each dataset size, we determined what percentage of these random ratios were greater than or equal to our actual observed ratios from the known O-GlcNAc proteins/ OGT interactors; this value was our permutation-test p-value.

#### Gene Set Enrichment Analysis of KEGG and Reactome Pathways

We downloaded all mouse pathways in the KEGG and Reactome databases (21, 22) and removed all genes not observed among our proteins, as well as pathways with fewer than 15 members, to provide 860 pathways tested. Protein list was truncated to remove isoforms; for each effect size, the isoform with the lowest p-value for that effect size was chosen, giving 7098 unique proteins, down from 7220 with multiple isoforms. Gene set enrichment analysis (GSEA) (23) was performed using pre-ranked settings for all 9 types of effect sizes (3 days x 3 OGT activities) as well as OGT inhibition data using mouse ortholog gene names (see below). To account for both statistical significance as well as effect size, GSEA was run twice for each effect size. One used effect sizes as the ranking metric, while the second used $-\log_{10}$ of p-value for effect size as the ranking metric; for genes with a negative effective size, $\log_{10}$ of p-value was used instead of $-\log_{10}$ to account for direction of change in response to the OGT activity. Benjamini-Hochberg-corrected false discovery rates (FDRs) were obtained for each set of pathways from both sets of ranking metrics, and hits were those meeting cutoffs of <1% FDR in one of the two GSEA analyses and at least <10% in the second one, allowing enrichment in a direction based on either highly-significant effect sizes or effect sizes representing large fold changes. The mean effect size (or $\log_2$ fold change) was obtained for all proteins in each pathway. If the sign of the mean did not match the direction of enrichment for a pathway, FDR was adjusted to 100%. 164 pathways met these cutoffs and are shown in Dataset S7. Selected pathways are shown in Figures 5B and S5D, with size of circle showing FDR based on $-\log_{10}$ p-value and color representing average pathway effect size or fold change.

### Comparison of Regression Coefficients to OGT Inhibition Data From Martin et al.

Data from Martin et al. 2018 (24) was downloaded, and BioMart (18) was used to find mouse orthologs of the HGNC symbols included in proteomics data from 293T samples. Multiday regression-derived coefficients were aligned to fold change data on these orthologous mouse gene IDs, yielding 5179 protein ortholog pairs present in both studies. $\mathrm{Log}_2$ of fold change was obtained for 10 $\mu$M OSMI4 treatment as was p-value in Welch's t-test of OGT inhibition versus DMSO treatment (n=2 per condition, comparison of $\log_2$ values). Data shown in Figures 5C and S5B was derived by obtaining all 513 proteins with orthologs in both studies that had $p<0.01$ for any effect size on any day. For each distribution in both figures, membership was determined based on direction of effect size after two days dTAG-13 treatment. p-values were based on wilcoxon ranked sum test of $\log_2$ fold change values. For correlation of pathways (Figure S5C), Pearson correlation coefficient was calculated between average $\log_2$ fold change for each pathway upon OGT inhibition and average coefficient fold change; this was performed for all 164 pathways included in Dataset S7, excluding those with <15 genes in the OGT inhibition dataset.

### Data and Code Availability

Raw mass spectrometry data for proteomics and imaging data from cell growth curves is available upon request. All custom code for data analysis and image production is available on GitHub at https://github.com/SuzanneWalkerLab/NoncatalyticOGT.

### Supplemental Figures

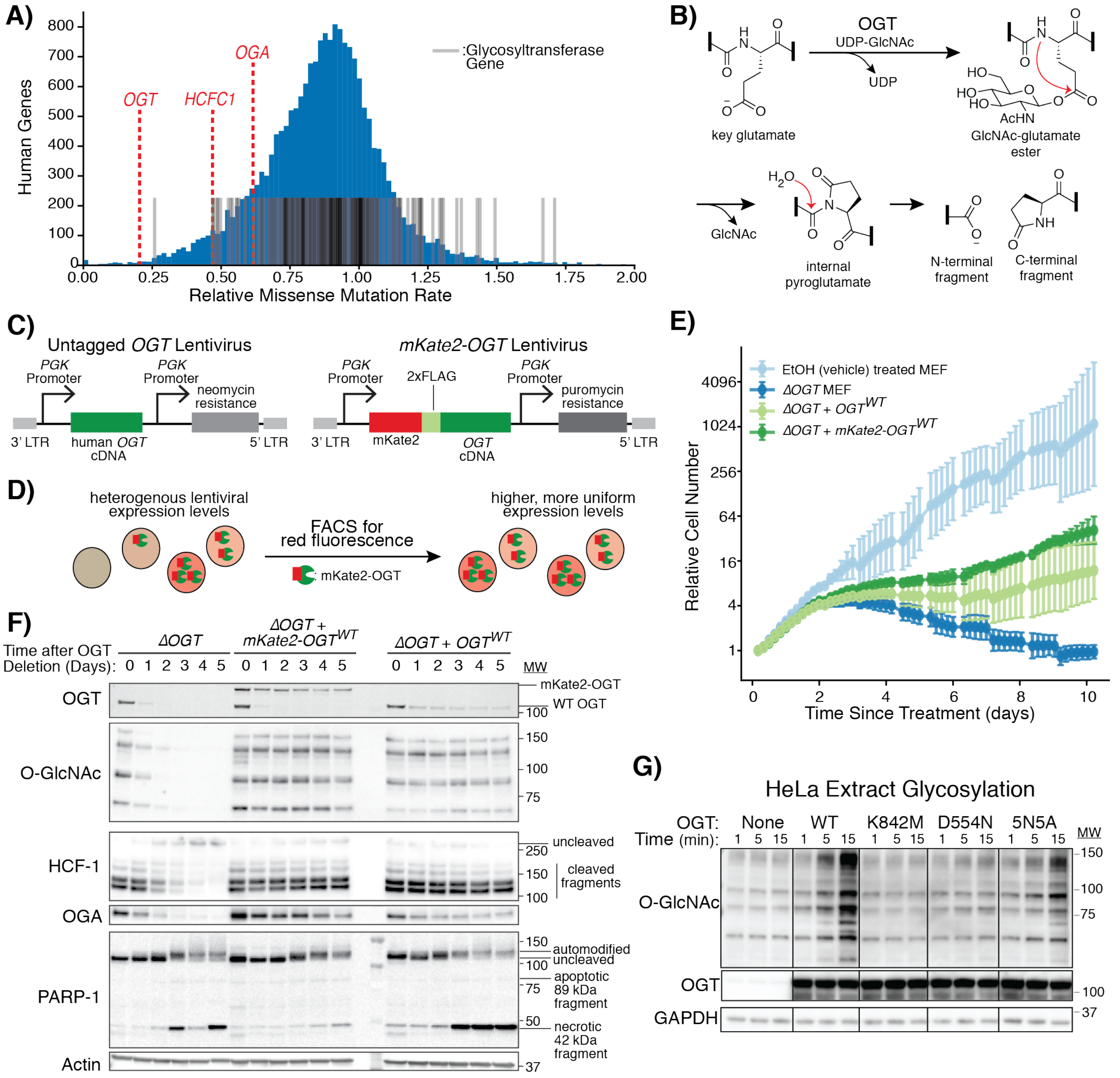


**Supplemental Figure 1: Figures Describing OGT Biochemical Activity and Strategies for Replacement of OGT in Cells (related to Figures 1–2)**

A) Histogram of human gene conservation from exome data as in Figure 1D (7), including *OGT, OGA*, and *HCFC1*, as well as marking other glycosyltransferase genes (8). Number of human genes plotted on y axis, with relative missense mutation rate (truncated at 2.0) on the x axis.

B) OGT glycosylates a glutamate on HCF-1 to form a GlcNAc-glutamate ester. Attack by the amide electron lone pair on the ester carbonyl produces an internal pyroglutamate. Water hydrolyzes this species to cleave the HCF-1 backbone. These reactions leave a pyroglutamate moiety on the C-terminal fragment. Vertical black bars show where the indicated residue joins the remainder of the HCF-1 protein.

C) Schematics of lentiviral constructs used to introduce *OGT*. The untagged *OGT* construct introduces the cDNA under the control of a *PGK* promoter alongside a neomycin resistance cassette. Constructs with *mKate2-OGT* have a 2xFLAG tag separating the fluorophore from OGT and are selectable by puromycin.

D) Schematic showing strategy to normalize OGT levels. Lentiviral *OGT* expression can be variable and low in many cells. After tagging OGT with the red fluorophore mKate2, we use fluorescence-activated cell sorting (FACS) to select a population of cells with robust *OGT* transgene expression.

E) Live cell imaging demonstrates rescue of growth with lentiviral *OGT* delivery. Cells were imaged every ~4 hours after *Ogt* knockout at t = 0. Doublings were measured by tracking confluence using the Incucyte Zoom live cell imaging system. Error bars are SD (n = 6 per condition).

F) Tagged OGT improves levels in cells without endogenous *Ogt*. Immunoblots track OGT biochemical activity in MEF cells after OGT removal without lentiviral infection, with mKate2-OGT, or with untagged OGT. Five days after *Ogt* removal, mKate2-OGT shows levels closer to endogenous OGT before genetic knockout, while untagged OGT shows lower levels. OGA levels are anticorrelated with OGT levels, while overall O-GlcNAc levels and degree of HCF-1 cleavage are constant when any *OGT* copy is present. Automodification of PARP-1, corresponding to DNA damage (25), is observed under all conditions, and no accumulation of apoptotic 89 kDa PARP-1 fragment is observed. Necrosis-associated 42 kDa fragment accumulates three days after OGT knockout in *ΔOGT* cells and those with untagged *OGT*, but not in *mKate2-OGT* cells (26).

G) Immunoblot analysis showing results of *in vitro* cell extract glycosylation experiments. Recombinant OGT variants glycosylated HeLa extracts for the indicated time at 37 °C. Reactions contained 2.5 μM recombinant OGT and 1 mM added UDP-GlcNAc. Pan-O-GlcNAc antibody CTD110.6 enabled detection of O-GlcNAc. 5N5A indicates mutant with mutation of 5 conserved Asn residues shown in Figure 2B (right) to Ala residues. For experiments in Figures 2–6, $OGT^{\Delta Cat}$ cells contains $\mathrm{OGT}^{\mathrm{K842M}}$, $OGT^{\Delta GlcNAc}$ cells contain $\mathrm{OGT}^{\mathrm{D554N}}$, and $OGT^{\Delta Clvg}$ cells contain $\mathrm{OGT}^{\mathrm{5N5A}}$.

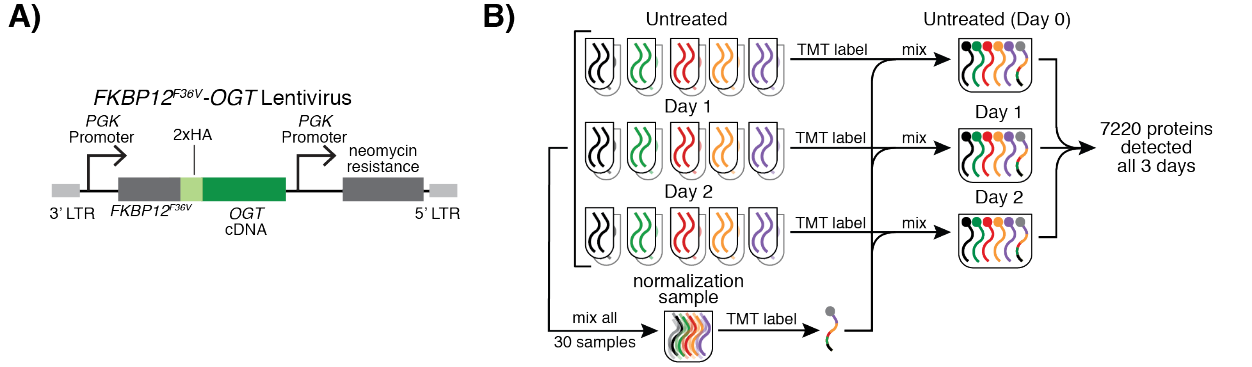


**Supplementary Figure 2: FKBP12$^{F36V}$-OGT Enables Proteomic Profiling of Effects of OGT Activities (Related to Figure 3)**

A) FKBP12$^{F36V}$-OGT consists of the FBKP12$^{F36V}$ variant fused via 2xHA linker. Neomycin-resistant lentivirus was used to insert the construct into *Ogt* knockout MEF cells.

B) A standardized sample was used to enable cross-day comparisons. We mixed aliquots of all 30 samples and labeled with an 11$^{th}$ TMT tag, which was added back to all 3 multiplexed proteomic runs. This channel was used for normalization between days.

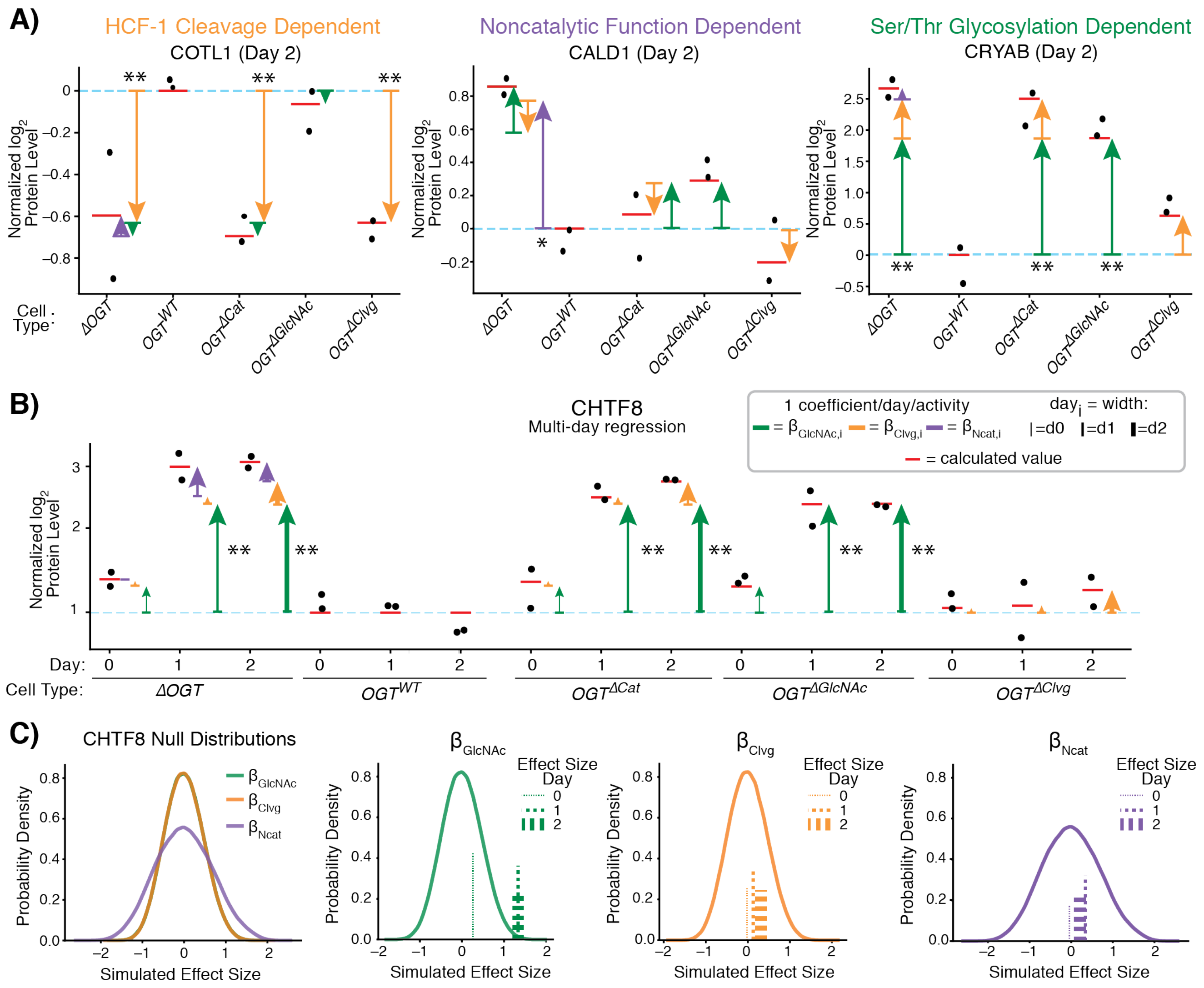


### Supplementary Figure 3: Linear Regression Measures Dependence of Protein Levels on Individual OGT Activities (Related to Figure 4)

A) Proteins primarily affected by a single OGT activity. For each protein, actual measured proteomic values after two days of dTAG-13 treatment are shown in black, calculated expression from regression coefficients is shown as red lines, and arrows indicate effect sizes of noncatalytic functions (purple), HCF-1 cleavage (yellow), and Ser/Thr glycosylation (green). Crystallin α B (CRYAB) protein levels are highly dependent on Ser/Thr glycosylation, coactosin like 1 (COTL1) protein levels are highly dependent on HCF-1 cleavage, and caldesmon 1 (CALD1) protein levels are highly dependent on noncatalytic functions. * $p < 0.05$; ** $p < 0.01$ (related to largest effect size only).

B) Multiday regression example. *$OGT^{WT}$* cells change limited amounts after dTAG-13 treatment, which allows us to measure changes from *$OGT^{WT}$* (i.e., loss of OGT activities) across all days. *$OGT^{WT}$* cells are used as the “constant” level for multiday regression. OGT-activity-driven changes in protein abundance for each day are calculated based on the degree of matching the “idealized” levels (as shown in Figure 4A). Separate coefficients are calculated for each day. CHTF8 is shown as an example protein with a large change in expression based on Ser/Thr glycosylation status. Samples are grouped by cell type, with the measured regression coefficients shown as vertical arrows colored by the corresponding OGT activity.

C) To obtain p-values, samples for each protein were bootstrap re-sampled and regression was repeated $10^6$ times. Actual measured effect sizes were compared to distributions from re-sampling, as shown for CHTF8. Similarities in data shape for each day’s effect sizes led to identical null distributions for each class of OGT activity. Due to limited number of samples lacking noncatalytic activities for each day (n = 2), higher variance is observed in null distributions for noncatalytic function effect sizes than for catalytic activities, leading to lower p-values for noncatalytic functions for the same effect size.

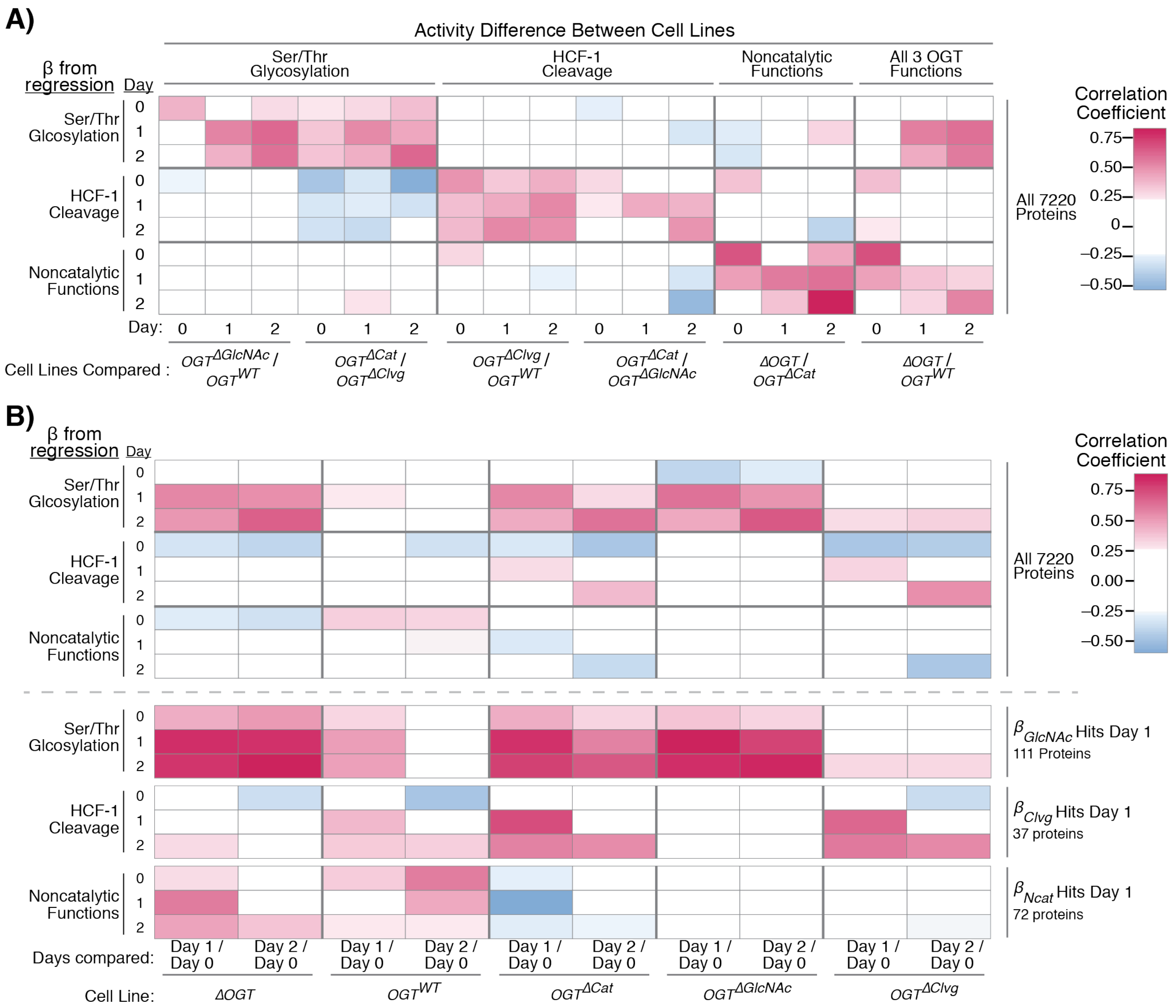


### Supplementary Figure 4: Correlation Analysis Validates Regression Approach (Related to Figure 4)

A) Heatmap of Pearson correlation coefficients between regression-derived effect sizes for each OGT activity at each timepoint (Y-axis) and $\log_2$ fold differences in protein abundance between pairs of cell lines (X-axis). Correlation is based on all 7220 proteins detected on all days. Cell line pairs are grouped based on the OGT activity they differ by (listed above heatmap). Cell line pairs either differ by a single OGT activity or by all three OGT activities simultaneously (*ΔOGT* v. $OGT^{WT}$). Protein abundance differences between cell line pairs differing by only one OGT activity positively correlated with the corresponding effect sizes for that OGT activity, especially when matched by timepoint. Protein differences between *ΔOGT* and $OGT^{WT}$ correlate with Ser/Thr glycosylation and noncatalytic functions, consistent with these two being OGT's dominant biochemical activities (Figure 4D)

B) Heatmap of Pearson correlation coefficients between regression-derived effect sizes (Y-axis) and $\log_2$ fold changes in protein abundance in single cell lines after 1 or 2 days of dTAG-13 treatment versus their untreated state (X-axis). Top panel shows correlation for all 7220 proteins, whereas bottom panels show correlations only for proteins scoring as hits for effect sizes for each OGT activity after 1 day of dTAG-13 treatment (effect size > $\log_2(1.5)$, $p < 0.05$). Correlation to each OGT activity in cell lines that lose that activity improves when only looking at Day 1 hit proteins, providing greater confidence that those proteins' levels are truly altered by the corresponding OGT activities.

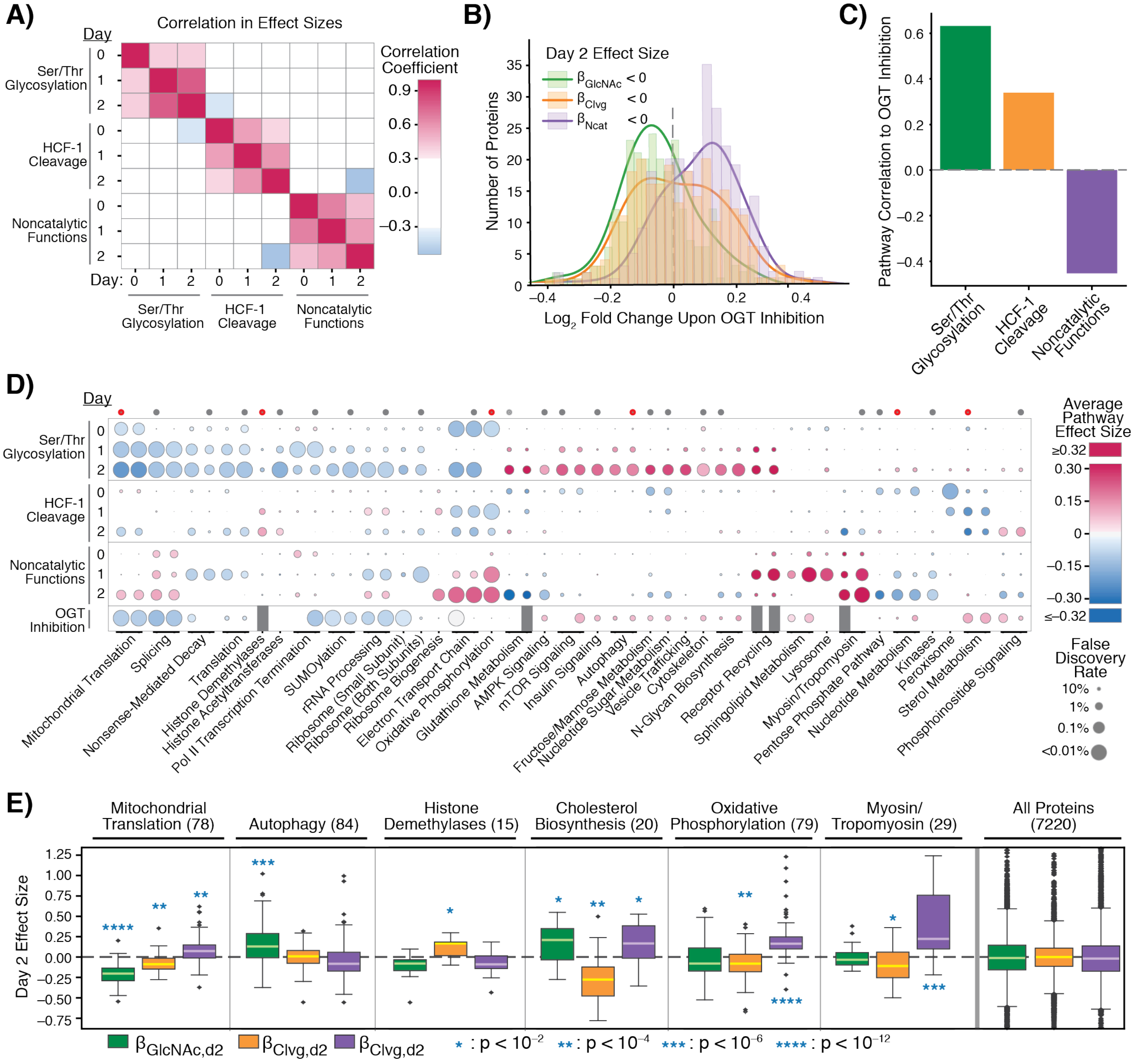


**Supplementary Figure 5: Proteins Dependent on Individual OGT Activities Differ From One Another (Related to Figure 5)**

A) Pearson correlation coefficients of effect sizes representing the three activities at all 3 days show limited correlation between three types of activities. All 7220 proteins included for correlation.

B) Related to Figure 5C. Histogram shows effect of OGT inhibition on protein levels for proteins with negative Day 2 $\beta_{GlcNAc}$ (n = 250), $\beta_{Clvg}$ (n = 249), and $\beta_{Ncat}$ (n = 278). Lines show smoothed distribution for each set of proteins (gaussian kernel with bandwidth of 0.05). Only proteins significant in regression analysis (p<0.01 for any effect size, 513 proteins) were analyzed; figure is sets of proteins with effect sizes in opposite direction of those displayed in Figure 5C. Based on Wilcoxon ranked sum test, only $\beta_{GlcNAc}$ and $\beta_{Ncat}$ distributions were more extreme than expected by chance (p-value $\beta_{GlcNAc} < 10^{-20}$, $\beta_{Clvg} \sim 0.14$, $\beta_{Ncat} < 10^{-13}$).

C) Pearson correlation between average pathway $\log_2$ fold changes upon OGT inhibition versus average effect sizes for all KEGG and Reactome pathways that were significantly associated with any OGT activity as described in Supplementary Methods.

D) GSEA shows pathways enriched for each OGT activity are different. GSEA was performed based on effect sizes from regression analysis and statistical significance; pathways shown are a selected subset of pathways meeting cutoffs for significant association. Color represents average effect size of all proteins in pathway for given OGT activity/day (Y-axis). OGT inhibition values are based on $\log_2$ fold change in protein abundance. Size of points is based on FDR from GSEA on basis of statistical significance. Pathways are listed based on categories; pathway names, average effect size, and FDRs based upon effect size or statistical significance available in Dataset S7. Dots above diagram indicate pathways in Figure 5B; red dots indicate pathways shown in *E).*

E) Selected pathways show trends in Day 2 effect sizes overall. Pathways indicated in *D)* are shown as box plots of all effect sizes for that pathway. Pathway category shown above each set of box plots, with number of proteins detected in pathway shown in parentheses. Asterisks indicate p-value comparing effect size to same category of Day 2 effect sizes for all proteins not in pathway in Wilcoxon ranked sum test. All proteins indicates effect sizes for all 7220 proteins measured in proteomics data; outliers continue beyond borders of plot for this sub-panel.

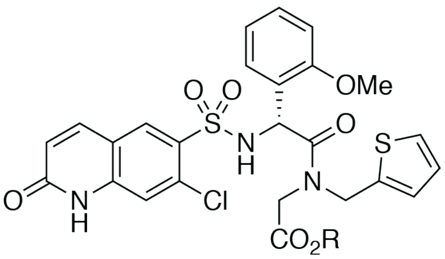


OGT Inhibitor OSMI-4

$K_d$ = 8 ± 1 nM

Cellular $EC_{50}$ = ~3 μM

**Supplementary Figure 6: OGT Inhibitor OSMI-4 Blocks OGT's Active Site (Related to Figure 5)**

OGT inhibitor OSMI-4 (24) potently binds OGT's active site, competing with UDP-GlcNAc. Compound is delivered as ester (R = Et) and thought to hydrolyze to acid in cells (R = H). $K_d$ for UDP-GlcNAc is 16 ± 4 μM. Cellular $EC_{50}$ based upon western blot analysis of global protein O-GlcNAcylation after 24 hours inhibitor treatment in HEK 293T cells. All comparisons in this study are to 24 hours treatment of HEK 293T cells with 10 μM OSMI-4.

### Supplementary Tables

*Table S1: Enrichment of Significant Effect Sizes in Known OGT Targets and Interactors*

| Category (# of proteins) | Glycosylated in Woo et al. (1250) | Binds OGT in Ruan et al. (314) |
|---|---|---|
| All Proteins (7220) | 17.3% of all measured proteins | 4.3% of all measured proteins |
| $\beta_{GlcNAc}$ $p < 0.01$ Day 2 (364) | 21.4% of proteins with significant $\beta_{GlcNAc}$ (enrichment $p < 0.02$) | 5.2% of proteins with significant $\beta_{GlcNAc}$ (enrichment $p > 0.2$) |
| $\beta_{Clvg}$ $p < 0.01$ Day 2 (91) | 23.1% of proteins with significant $\beta_{Clvg}$ (enrichment $p > 0.05$) | 5.5% of proteins with significant $\beta_{Clvg}$ (enrichment $p > 0.7$) |
| $B_{Ncat}$ $p < 0.01$ Day 2 (76)* | 17.1% of proteins with significant $\beta_{Ncat}$ (enrichment $p > 0.5$) | 10.5% of proteins with significant $\beta_{Ncat}$ (enrichment $p < 0.02$) |

*Note that the higher variance of noncatalytic function effect sizes leads to increased p-values relative to catalytic activities.

Data for glycosylated targets and interactors from references (25) and (26). p-values obtained using permutation test.

*Table S2: Plasmids Used or Generated in Study*

| Plasmid Name | Plasmid Source | Addgene # |
|---|---|---|
| pENTR1A (no ccdB) | Campeau et al.(27) | 17398 |
| pLenti PGK Neo DEST | Campeau et al.(27) | 19067 |
| pLenti PGK Puro DEST | Campeau et al. 2009(27) | 19068 |
| pLEX_305-N-dTAG | Nabet et al. 2018 (28) | 91797 |
| WT OGT Bacterial Expression Plasmid | Gross et al. 2006 (29) | 154271 |
| K842M OGT Bacterial Expression Plasmid | Lazarus et al. 2013 (30) | 154272 |
| D554N OGT Bacterial Expression Plasmid | Janetzko et al. 2016 (31) | 154279 |
| 5N5A OGT Bacterial Expression Plasmid | Levine et al. 2018 (2) | 154280 |
| Human WT OGT pENTR1A | This paper | 154281 |
| Human WT mKate2-OGT pENTR1A | This paper | 154282 |
| Human 5N5A mKate2-OGT pENTR1A | This paper | 154285 |
| Human D554N mKate2-OGT pENTR1A | This paper | 154283 |
| Human K842M mKate2-OGT pENTR1A | This paper | 154284 |

| | | |
|---|---|---|
| Human WT OGT pLenti PGK Neo | This paper | 154289 |
| Human WT mKate2-OGT pLenti PGK Puro | This paper | 154290 |
| Human D554N mKate2-OGT pLenti PGK Puro | This paper | 154292 |
| Human 5N5A mKate2-OGT pLenti PGK Puro | This paper | 154293 |
| Human K842M mKate2-OGT pLenti PGK Puro | This paper | 154291 |
| Human WT OGT N-term open pENTR1A | This paper | 155197 |
| FKBP12$^{F36V}$-OGT pENTR1A | This paper | 154287 |
| FKBP12$^{F36V}$-OGT pLenti PGK Neo | This paper | 154294 |
| pLEX_305-N-dTAG_OGT | This paper | 155196 |
| psPAX2 | gift from Didier Trono | 12260 |
| pMD2.G | gift from Didier Trono | 12259 |

*Table S3: Primers and dsDNA Fragments Used in Study*

| Single-Stranded Oligonucleotides | |
|---|---|
| **Name/Description** | **Sequence** |
| Human OGT cDNA Amplification 5' FWD BamHI | AAGGAACCAATTCAGTCGACTGGATCCATGGCGTCT TCCGTGGG |
| Human OGT cDNA Amplification 3' REV XhoI | ACTTTGTACAAGAAAGCTGGGTCTAGATATCTCGAG TTATGCTGACTCAGTGACTTCAAC |
| Human OGT K842M mutation FWD | TCAGTTGTATatgATTGACCCTTCTAC |
| Human OGT K842M mutation REV | TTAAAGTTACAGTATACGATG |
| Human OGT D554N Mutation FWD | TGTGAGTTCCaacTTTGGGAATCATC |
| Human OGT D554N Mutation REV | TATCCTACACGCAGCCGA |
| Human OGT linearization for 5N5A mutation FWD | GCCTGCAGATTGTCTGTGATT G |
| Human OGT linearization for 5N5A mutation REV | TCTGCATGGGTGGGACAC |
| OGT pENTR1A linearization for mKate2-OGT | CATGGTGGCGGCCGACTG |

| | |
|---|---|
| FWD | |
| OGT pENTR1A linearization for mKate2-OGT REV | GCGTCTTCCGTGGGCAAC |
| OGT “open” N-terminus pENTR1A FWD | GCGTCTTCCGTGGGCAAC |
| OGT “open” N-terminus pENTR1A REV (BamHI site) | AAAGCCTGCTTTTTTGTACAAAGTTG |
| FKBP12$^{F36V}$-OGT PCR FWD (Not | ATATATGGATCCATGGGAGTGCAGGTGGAAACCATC |
| FKBP12$^{F36V}$-OGT PCR REV | ATATATGCGGCCGCGTTATGCTGACTCAGTGACTTC<br>AACAGGC |
| **Double Stranded DNA** | |
| mKate2-2xFLAG gBlock | |
| GTCGGCCGCCACCATGGTGAGCGAGCTGATTAAGGAGAACATGCACATGAAGCTGTACATGGAGGGCACC<br>GTGAACAACCACCACTTCAAGTGCACATCCGAGGGCGAAGGCAAGCCCTACGAGGGCACCCAGACCATGA<br>GAATCAAGGCGGTCGAGGGCGGCCCTCTCCCCTTCGCCTTCGACATCCTGGCTACCAGCTTCATGTACGG<br>CAGCAAAACCTTCATCAACCACACCCAGGGCATCCCCGACTTCTTTAAGCAGTCCTTCCCCGAGGGCTTCA<br>CATGGGAGAGAGTCACCACATACGAAGACGGGGGCGTGCTGACCGCTACCCAGGACACCAGCCTCCAGG<br>ACGGCTGCCTCATCTACAACGTCAAGATCAGAGGGGTGAACTTCCCATCCAACGGCCCTGTGATGCAGAAG<br>AAAACACTCGGCTGGGAGGCCTCCACCGAGACCCTGTACCCCGCTGACGGCGGCCTGGAAGGCAGAGCC<br>GACATGGCCCTGAAGCTCGTGGGCGGGGGCCACCTGATCTGCAACTTGAAGACCACATACAGATCCAAGA<br>AACCCGCTAAGAACCTCAAGATGCCCGGCGTCTACTATGTGGACAGAAGACTGGAAAGAATCAAGGAGGC<br>CGACAAAGAGACCTACGTCGAGCAGCACGAGGTGGCTGTGGCCAGATACTGCGACCTCCCTAGCAAACTG<br>GGGCACAGAGGCGGAGACTACAAAGATGATGACGACAAAGGCGACTACAAAGACGATGACGACAAGGGCA<br>TCGATAGATCAACAAGTTTGTACAAAAAAGCAGGCTTTGCGTCTTCCGTGGGCA | |
| gBlock for 5N5A mutation human OGT (capitalized letters indicate sites of mutated residues) | |
| cccacccatgcagactctctgaatGCActagccaatatcaaacgagaacagggaaacattgaagaggcagttcgcttgtatcgtaaagcattagaagtctt<br>cccagagtttgctgctgcccattcaGCAttagcaagtgtactgcagcagcagggaaaactgcaggaagctctgatgcattataaggaggctattcgaatcag<br>tcctacctttgctgatgcctactctGCAatgggaaacactctaaaggagatgcaggatgttcagggagccttgcagtgttatacgcgtgccatccaaattaatc<br>ctgcatttgcagatgcacatagcGCActggcttccattcataaggattcagggaatattccagaagccatagcttcttaccgcacggctctgaaacttaagcct<br>gattttcctgatgcttattgtGCAttggctcattgcctgcagattgtctg | |

### Supplementary Datasets

*Dataset S1: Relative Expression of All Proteins Quantified By Quantified Mass Spectrometry (Before Normalization)*

Includes Protein Id, Gene Symbol, and text description of all proteins identified by mass spectrometry, along with peptide counts from each single day TMT proteomics experiment. Total relative quantified abundance for each cell type for each protein is listed, with 0 indicating protein not detected by mass spectrometry for that time point. Note “clone” refers to different FKBP12$^{F36V}$-OGT clonal cell background, serving as different biological replicates for each cell line, and normalization standard refers to sample made of a mix of all 30 samples that is used in mass spectrometry runs for each timepoint to enable inter-day normalization.

*Dataset S2: Normalized Levels of Proteins Across Multiple Days*

Table of $\log_2$ mean expression normalized levels of all proteins detected with ≥1 peptide at all 3 days. Also includes $\log_2$ fold difference and 2 sample t-test p-value (on $\log_2$ normalized expression) for Day 0 versus Day 1 or Day 2 of dTAG-13 treatment within the same cell line. Hits for each inter-day comparison are also listed as True (for hits with > 1.5 fold change in either direction and a two-sided t-test p-value < 0.01) or False. UniProt Accession, Protein Id, Gene Symbol, and Description given as protein identifiers.

*Dataset S3: Proteome-Wide Effects of OGT Activities on Protein Levels Day 2 dTAG-13 Treatment*

Table of linear regression-derived effect sizes describing association between individual OGT biochemical activities and proteome-wide protein level changes, as described in Figure 4B. Betas are effect sizes that represent $\log_2$ fold effect of loss of activity on protein level for each OGT activity (GlcNAc = Ser/Thr Glycosylation, Clvg = HCF-1 Cleavage, Ncat = Noncatalytic functions). Associated p-value describes likelihood of obtaining actual effect size if there was no association between cell type and protein level. Baseline expression from $OGT^{WT}$ cell line also shown.

*Dataset S4: Proteome-Wide Effects of OGT Activities on Protein Levels (Multi-Day Regression)*

Table of multi-day regression-derived effect sizes for each OGT activity at each day (as described in Supplementary Figure 3B). Betas are effect sizes in $\log_2$ fold difference for loss of activity for that day. Associated p-value describes likelihood of obtaining effect size if there was no association between cell type and protein level. Hits for a given activity-day pair are True if they have >1.5 fold change in either direction for that activity on that day with $p < 0.05$. Baseline expression *C* (average $OGT^{WT}$ level of that protein) is also given with p-value for likelihood of $OGT^{WT}$ cells having a stable protein level relative to protein abundance differences between cell lines. Includes whether protein is glycosylated in Woo et al.(19) or binds OGT as measured by Ruan et al.(20)

*Dataset S5: Proteome-Wide Single-Day Comparisons of Protein Abundance Between Cell Types*

Table of $\log_2$ fold changes for differences in abundance of given proteins in same-day comparisons between cell line pairs shown in Figure S4A, along with associated p-values from Welch's t-test on $\log_2$ transformed data.

*Dataset S6: Protein Ortholog Abundance Changes from OGT Inhibition*

Protein $\log_2$ fold change in abundance (and t-test p-value) in HEK 293T cells at 24 hours in response to OGT inhibition (24). Both mouse and human gene name given for each protein ortholog to facilitate comparison to effect sizes from mouse genes in MEFs.

*Dataset S7: Pathways Altered By OGT Activities and OGT Inhibition*

GSEA pathway analysis showing Reactome and KEGG pathways. GSEA was run on both degree of statistical significance (as measured by $-\log_{10}$ of p-value) and effect sizes- 169 pathways met cutoffs of <10% FDR in one of the two GSEA methods and <1% FDR in the other. Also listed are degrees of enrichment for similar anlyses conducted on $\log_2$ protein abundance changes due to OGT inhibition in HEK cells. Order is same as in Figures 5B, S5D, but with complete list of pathways (pathways in figures are indicated).

### Supplementary References


1. Z. Kazemi, H. Chang, S. Haserodt, C. McKen, N. E. Zachara, O-Linked β-N-acetylglucosamine (O-GlcNAc) Regulates Stress-induced Heat Shock Protein Expression in a GSK-3β-dependent Manner. *J. Biol. Chem.* **285**, 39096–39107 (2010).
2. Z. G. Levine, *et al.*, O-GlcNAc Transferase Recognizes Protein Substrates Using an Asparagine Ladder in the Tetratricopeptide Repeat (TPR) Superhelix. *J. Am. Chem. Soc.* **140**, 3510–3513 (2018).
3. D. G. Gibson, Synthesis of DNA fragments in yeast by one-step assembly of overlapping oligonucleotides. *Nucleic Acids Res.* **37**, 6984–6990 (2009).
4. J. Navarrete-Perea, Q. Yu, S. P. Gygi, J. A. Paulo, Streamlined Tandem Mass Tag (SL-TMT) Protocol: An Efficient Strategy for Quantitative (Phospho)proteome Profiling Using Tandem Mass Tag-Synchronous Precursor Selection-MS3. *J. Proteome Res.* **17**, 2226–2236 (2018).
5. J. A. Paulo, J. D. O'Connell, S. P. Gygi, A Triple Knockout (TKO) Proteomics Standard for Diagnosing Ion Interference in Isobaric Labeling Experiments. *J. Am. Soc. Mass Spectrom.* **27**, 1620–1625 (2016).
6. G. C. McAlister, *et al.*, MultiNotch MS3 enables accurate, sensitive, and multiplexed detection of differential expression across cancer cell line proteomes. *Anal. Chem.* **86**, 7150–7158 (2014).
7. K. J. Karczewski, *et al.*, The mutational constraint spectrum quantified from variation in 141,456 humans. *Nature* **581**, 434–443 (2020).

8. M. Jöud, M. Möller, M. L. Olsson, Identification of human glycosyltransferase genes expressed in erythroid cells predicts potential carbohydrate blood group loci. *Sci. Rep.* **8**, 6040 (2018).
9. J. A. Paulo, S. P. Gygi, Nicotine-induced protein expression profiling reveals mutually altered proteins across four human cell lines. *Proteomics* **17** (2017).
10. J. A. Paulo, S. P. Gygi, Isobaric Tag-Based Protein Profiling of a Nicotine-Treated Alpha7 Nicotinic Receptor-Null Human Haploid Cell Line. *Proteomics* **18**, e1700475 (2018).
11. J. A. Paulo, S. P. Gygi, mTMT: An Alternative, Nonisobaric, Tandem Mass Tag Allowing for Precursor-Based Quantification. *Anal. Chem.* **91**, 12167–12172 (2019).
12. S. A. Beausoleil, J. Villén, S. A. Gerber, J. Rush, S. P. Gygi, A probability-based approach for high-throughput protein phosphorylation analysis and site localization. *Nat. Biotechnol.* **24**, 1285–1292 (2006).
13. E. L. Huttlin, *et al.*, A tissue-specific atlas of mouse protein phosphorylation and expression. *Cell* **143**, 1174–1189 (2010).
14. J. E. Elias, S. P. Gygi, Target-decoy search strategy for increased confidence in large-scale protein identifications by mass spectrometry. *Nat. Methods* **4**, 207–214 (2007).
15. J. E. Elias, S. P. Gygi, Target-decoy search strategy for mass spectrometry-based proteomics. *Methods Mol. Biol. Clifton NJ* **604**, 55–71 (2010).
16. G. C. McAlister, *et al.*, Increasing the multiplexing capacity of TMT using reporter ion isotopologues with isobaric masses. *Anal. Chem.* **84**, 7469–7478 (2012).
17. J. H. Jr. Ward, Hierarchical Grouping to Optimize an Objective Function. *J. Am. Stat. Assoc.* **58**, 236–244 (1963).
18. D. Smedley, *et al.*, BioMart – biological queries made easy. *BMC Genomics* **10**, 22 (2009).
19. C. M. Woo, *et al.*, Mapping and Quantification of Over 2000 O-linked Glycopeptides in Activated Human T Cells with Isotope-Targeted Glycoproteomics (Isotag). *Mol. Cell. Proteomics MCP* **17**, 764–775 (2018).
20. H.-B. Ruan, *et al.*, O-GlcNAc transferase/host cell factor C1 complex regulates gluconeogenesis by modulating PGC-1α stability. *Cell Metab.* **16**, 226–237 (2012).
21. B. Jassal, *et al.*, The reactome pathway knowledgebase. *Nucleic Acids Res.* **48**, D498–D503 (2020).
22. M. Kanehisa, S. Goto, KEGG: kyoto encyclopedia of genes and genomes. *Nucleic Acids Res.* **28**, 27–30 (2000).
23. A. Subramanian, *et al.*, Gene set enrichment analysis: A knowledge-based approach for interpreting genome-wide expression profiles. *Proc. Natl. Acad. Sci.* **102**, 15545–15550 (2005).
24. S. E. S. Martin, *et al.*, Structure-Based Evolution of Low Nanomolar O-GlcNAc Transferase Inhibitors. *J. Am. Chem. Soc.* **140**, 13542–13545 (2018).
25. G. V. Chaitanya, J. S. Alexander, P. P. Babu, PARP-1 cleavage fragments: signatures of cell-death proteases in neurodegeneration. *Cell Commun. Signal. CCS* **8**, 31 (2010).
26. S. Gobeil, C. C. Boucher, D. Nadeau, G. G. Poirier, Characterization of the necrotic cleavage of poly(ADP-ribose) polymerase (PARP-1): implication of lysosomal proteases. *Cell Death Differ.* **8**, 588–594 (2001).
27. E. Campeau, *et al.*, A Versatile Viral System for Expression and Depletion of Proteins in Mammalian Cells. *PLoS ONE* **4**, e6529 (2009).
28. B. Nabet, *et al.*, The dTAG system for immediate and target-specific protein degradation. *Nat. Chem. Biol.* **14**, 431 (2018).
29. B. J. Gross, B. C. Kraybill, S. Walker, Discovery of O-GlcNAc transferase inhibitors. *J. Am. Chem. Soc.* **127**, 14588–14589 (2005).

30. M. B. Lazarus, *et al.*, HCF-1 is cleaved in the active site of O-GlcNAc transferase. *Science* **342**, 1235–1239 (2013).
31. J. Janetzko, S. A. Trauger, M. B. Lazarus, S. Walker, How the glycosyltransferase OGT catalyzes amide bond cleavage. *Nat. Chem. Biol.* **12**, 899–901 (2016).
